# Comammox *Nitrospira* are the dominant ammonia oxidizers in a mainstream low dissolved oxygen nitrification reactor

**DOI:** 10.1101/504704

**Authors:** Paul Roots, Yubo Wang, Alex F. Rosenthal, James S. Griffin, Fabrizio Sabba, Morgan Petrovich, Fenghua Yang, Joseph A. Kozak, Heng Zhang, George F. Wells

**Affiliations:** Department of Civil and Environmental Engineering, Northwestern University, 2145 Sheridan Road, Evanston, IL, 60208, USA; Metropolitan Water Reclamation District of Greater Chicago, 6001 W Pershing Road, Chicago,IL, 60804, USA

**Keywords:** Biological nutrient removal (BNR), energy efficient, wastewater, nitrogen cycling

## Abstract

Recent findings show that a subset of bacteria affiliated with *Nitrospira*, a genus known for its importance in nitrite oxidation for biological nutrient removal applications, are capable of <u>co</u>mplete <u>amm</u>onia <u>ox</u>idation (comammox) to nitrate. Early reports suggested that they were absent or present in low abundance in most activated sludge processes, and thus likely functionally irrelevant. Here we show the accumulation of comammox *Nitrospira* in a nitrifying sequencing batch reactor operated at low dissolved oxygen (DO) concentrations. Actual mainstream wastewater was used as influent after primary settling and an upstream pre-treatment process for carbon and phosphorus removal. The ammonia removal rate was stable and exceeded that of the treatment plant’s parallel full-scale high DO nitrifying activated sludge reactor. 16S rRNA sequencing showed a steady accumulation of *Nitrospira* to 53% total abundance and a decline in conventional ammonia oxidizing bacteria to <1% total abundance over 400+ days of operation. After ruling out other known ammonia oxidizers, qPCR confirmed the accumulation of comammox *Nitrospira* beginning around day 200, to eventually comprise 94% of all detected *amoA* and 4% of total bacteria by day 407. Quantitative fluorescence in-situ hybridization confirmed the increasing trend and high relative abundance of *Nitrospira*. These results demonstrate that comammox can be metabolically relevant to nitrogen transformation in wastewater treatment, and can even dominate the ammonia oxidizing community. Our results suggest that comammox may be an important functional group in energy efficient nitrification systems designed to operate at low DO levels.

## 1. Introduction

Resource and energy-efficient nutrient removal methods from wastewater have gained increased attention in recent years due to utility goals of energy self-sufficiency and increasingly stringent effluent nitrogen (N) and phosphorus (P) standards. Such methods include partial nitritation/anammox (PN/A) and nitritation-denitritation, both of which require low dissolved oxygen (DO) or anoxic zones, as well as simply nitrification with low DO. These methods contrast with conventional nitrifying activated sludge operated at high, i.e. >2 mg/L DO concentrations (Park and Noguera, 2004), which incur substantial energy costs due to aeration. When low DO methods are applied to mainstream wastewater, they may select for organisms that thrive under relatively low substrate (i.e. due to stringent effluent standards) and low oxygen conditions.

Comammox *Nitrospira*, discovered in late 2015 (Daims et al., 2015; van Kessel et al., 2015), may be such an organism. The name “comammox” is derived from their ability to perform <u>com</u>plete <u>amm</u>onia <u>ox</u>idation all the way to nitrate (NO_3_^−^). The comammox metabolism overturned a >100-year paradigm that nitrification is a two-step process requiring coordinated activity of ammonia oxidizing bacteria or archaea (AOB or AOA, ammonium [NH_4_^+^] to nitrite [NO_2_^−^]) and nitrite oxidizing bacteria (NOB, NO_2_^−^ to NO_3_^−^). The presence of comammox could be detrimental to PN/A, nitritation, and nitritation-denitritation process performance, where accumulation of the intermediate NO_2_^−^ via NOB suppression is a process requirement, and NO_3_^−^ production is not desired. Early mining of shotgun metagenomics datasets revealed their presence (albeit low abundance) in conventional nitrifying activated sludge systems (Daims et al., 2015; van Kessel et al., 2015), as their unique ammonia monooxygenase (*amoA*) gene had defied detection by previous quantitative polymerase chain reaction (qPCR) or sequencing assays targeting AOB. The extent of their importance to wastewater treatment, however, is currently unknown.

As demonstrated by substrate affinity tests on axenic cultures of *Nitrospira inopinata* (Kits et al., 2017), at least one comammox species is adapted to oligotrophic conditions due to its very high NH_4_^+^ affinity (half-saturation coefficient K_NH4+_ = 0.01 mgNH_4_^+^-N/L) and relatively low maximum rate of ammonium oxidation. While the oxygen affinity of comammox has yet to be measured, theoretical predictions and genomic studies alike indicate that they are likely adapted to environments with low DO (Costa et al., 2006; Lawson and Lücker, 2018). These characteristics are borne out in the conditions in which comammox has been discovered to date. Indeed, *N. inopinata* was originally cultured from a biofilm growing 1,200 m below the surface in an oil exploration well (Daims et al., 2015), while *Candidatus* Nitrospira nitrosa and *Candidatus* Nitrospira nitrificans were first found in biomass on an aquaculture system biofilter (van Kessel et al., 2015) exposed to NH_4_^+^ concentrations of 1.1 mgNH_4_^+^-N/L or less. Comammox *Nitrospira* were also found in an aquaculture biofilter with low (0.1 mgNH_4_^+^-N/L) substrate concentrations (Bartelme et al., 2017), and in this case outnumbered both AOA and AOB based on *amoA* quantification via qPCR. Subsequent investigations found comammox in the oligotrophic environment of drinking water treatment plants (Fowler et al., 2018; Palomo et al., 2016; Pinto et al., 2016) where they were detected in 10 of 12 locations with metagenomic datasets (Wang et al., 2017). In four of those samples, comammox *amoA* outnumbered that of AOA and AOB, suggesting that comammox may dominate nitrification activity in some drinking water treatment plants.

The “oligotrophic lifestyle” (Kits et al., 2017) of comammox suggests that they may not be able to compete in the relatively nutrient-rich environments of wastewater treatment plants (WWTPs), and two surveys for comammox in WWTPs seem to confirm this (Annavajhala et al., 2018; Gonzalez-Martinez et al., 2016). Gonzalez-Martinez et al. (2016) utilized 16S rRNA sequencing to survey 6 full-scale nitrifying activated sludge systems and 3 full-scale autotrophic nitrogen removal systems and found only one operational taxonomic unit (OTU) affiliated with comammox *Nitrospira* at a very low abundance of 0.08%, or five times less abundant than AOB. Annavajhala et al. (2018) examined metagenomic data sets of 16 full-scale biological nitrogen removal systems and found comammox *Nitrospira* present in all reactors at a relative abundance of 0.28 – 0.64% (Annavajhala et al., 2018). All samples had a ratio of comammox to AOB protein coding sequences of 0.18 or less, suggesting that comammox played a relatively minor role in NH_4_^+^ oxidation in all systems. While the conclusion of Gonzalez-Martinez et al. (2016) was that comammox are not significant in nitrogen cycling, Annavajhala et al. (2018) cautioned that their seeming ubiquity in WWTPs suggests that further research is warranted to understand their contribution to nitrogen transformations.

Increased research into low DO N removal systems for treatment of mainstream wastewater (as opposed to sidestream systems with high N concentrations > 300 mgN/L) to reduce energy consumption by aeration may produce environments more favorable to the oligotrophic comammox. Here, we demonstrate strong enrichment of comammox coupled to high rate complete ammonia oxidation activity in a low DO nitrifying SBR operated for >400 days with real primary effluent as feed. While comammox *Nitrospira* have been previously detected in wastewater treatment systems, this study is the first to show it to be the dominant ammonia oxidizer in a mainstream wastewater treatment bioreactor without synthetic feed.

## 2. Methods

### 2.1 SBR operation/control, inoculation, online sensors, and batch activity assays

A 12-L suspended growth sequencing batch reactor (SBR) was fed pre-treated primary effluent at the Metropolitan Water Reclamation District of Greater Chicago Terrance J. O’Brien Water Reclamation Plant (WRP) in Skokie, Illinois, USA for 414 days. The SBR included online sensors (s::can, Vienna, Austria) for monitoring of temperature, DO, ammonium and pH every minute. The reactor was initially operated with the intent to select for mainstream suspended growth PN/A, but was transitioned over the course of reactor operation to a low DO complete nitrification reactor. Upstream treatment included primary settling tanks and a 56-L A-stage activated sludge sequencing batch reactor for COD and biological phosphorus removal. The 12-L reactor was seeded on May 24, 2016 (day 0) with ∼1,800 mg/L of mixed liquor volatile suspended solids (MLVSS) suspended growth biomass from the full-scale sidestream DEMON® process at the York River treatment plant (Hampton Roads Sanitation District; HRSD) and ∼200 mgVSS/L of scraped biofilm from the Kruger/Veolia Biofarm (ANITA™ Mox process) at James River, VA (equivalent to 10% of the initial VSS). The reactor was subsequently loaded with A-stage effluent (Table 1), temperature controlled to 20.3 ± 1.1°C, and operated as a SBR to approximate plug flow reactor (PFR) behavior. Over the entire study, the average MLVSS was 1.4 ± 0.4 g VSS/L and solids were not intentionally wasted from the reactor, resulting in an average SRT of 99 ± 44 days (see Supporting Information for details on the SRT calculation).

**Table 1.** Low DO nitrification reactor influent (A-stage effluent) and effluent average composition, along with primary effluent and parallel full-scale (O’Brien WRP) nitrifying activated sludge bioreactor effluent concentrations.

|  | Primary effluent | A-stage effluent | Reactor effluent | O'Brien effluent |
| --- | --- | --- | --- | --- |
| TKN (mgN/L) | 20.6 ± 4.4 | 16.5 ± 4.7 | 4.5 ± 2.7 | 1.9 ± 0.2 |
| NH<sub>4</sub><sup>+</sup> (mgN/L) | 15.5 ± 3.6 | 14.3 ± 3.8 | 3.6 ± 2.6 | 0.7 ± 0.1 |
| NO<sub>x</sub> (mgN/L) | ---<sup>a</sup> ± --- | 0.5<sup>b</sup> ± 0.7 | 7.2 ± 3.3 | 7.4 ± 2.1 |
| COD (mgCOD/L) | 141 ± 43 | 42 ± 32 | 24 ± 17 | not available<sup>c</sup> |
| sCOD (mgCOD/L) | 84 ± 21 | 29 ± 11 | 20 ± 7 | not available |
| alkalinity (meq/L) | 4.7 ± 0.5 | 4.6 ± 0.5 | 3.3 ± 0.6 | not available |
| TSS (mg/L) | 45 ± 25 | 15 ± 35 | 7 ± 8 | 6 |

<sup>a</sup>NO<sub>x</sub> in primary effluent was at or below detection limit of 0.15 mgN/L in 93% of samples
<sup>b</sup>NO<sub>x</sub> in A-stage effluent was at or below detection limit of 0.15 mgN/L in 54% of samples
<sup>c</sup>COD not measured, but BOD<sub>5</sub> in O'Brien WRP effluent = 5.7 ± 2.9 mgBOD/L

SBR control of reactor equipment from inoculation to day 336 was managed with on-off circuit switching via ChronTrol programmable timers (4-circuit, 8-input XT Table Top unit, ChronTrol, San Diego, CA, USA). Sequences consisted of an initial fill + anaerobic react phase (4 - 30 minutes including 4-minute fill), an intermittently aerated react phase (240 - 270 minutes), settling (30 minutes), and decant (5 minutes). Aeration intervals were varied throughout the project depending on influent strength and aeration strategy from 1 – 2 minutes of aeration in a 2 – 30-minute interval. Target peak DO concentrations during aeration varied between 0.2 – 1.0 mgO_2_/L. Not including settling, the 50% volume decant resulted in a 9-hour hydraulic residence time (HRT).

Starting on day 337 and continuing to the end of the study (day 414), reactor equipment was controlled with code-based Programmable Logic Control (PLC) (Ignition SCADA software by Inductive Automation, Fulsom, CA, USA, and TwinCAT PLC software by Beckhoff, Verl, Germany). Aeration control was switched to proportional-integral (PI) control based on the online oxygen sensor (s::can oxi::lyser™ optical probe) signal. On day 358, the length of the aerated react portion of the cycle switched from timer-based to ammonium sensor-based (s::can ammo::lyser™ ion selective electrode) control. The aerated react period ended when NH_4_^+^ dropped to 2 mgNH_4_^+^ - N/L, resulting in variable HRT. Cycle phases from day 358 to day 414, the end of the study, consisted of fill (∼2 minutes), anaerobic react (20 minutes), intermittently aerated react (average 112 ± 69 minutes), polishing anaerobic react (20 minutes), settling (30 minutes), and decant (4.5 minutes). Not including settling and decant/fill, this resulted in a 5.1 ± 2.3-hour HRT.

Batch kinetic assays were performed to determine maximum activities of anammox, ammonia-oxidizing microorganisms (AOM), and NOB functional groups under non-limiting substrate conditions, as previously described (Laureni et al., 2016). See Supporting Information for details.

### 2.2 Full-scale secondary treatment at O’Brien WRP

The bench-scale SBR described above for low DO nitrification was operated in parallel to a full-scale secondary treatment process at the O’Brien WRP consisting of one-stage conventional activated sludge in plug-flow configuration (O’Brien) targeting biochemical oxygen demand (BOD) removal and NH_4_^+^ oxidation (nitrification). O’Brien received the same influent, or primary effluent (Table 1), as the A-stage to the bench-scale SBR (though in continuous feed mode vs. batch), offering a comparison point for the nitrifying community selected under similar influent but differing operating conditions. The O’Brien system was operated with an approximate average HRT of 7.3 hours, an SRT of 9.7 days, and MLVSS of 1.9 g/L. Wastewater temperature varied seasonally (low monthly average = 11°C, high monthly average = 22°C) with an average of 16.9°C. Aeration was provided throughout the basins (in contrast to intermittent aeration in the bench-scale reactor), and DO near the end of the basins was typically between 3 – 5 mgO_2_/L (in contrast to 0.2 – 1 mgO_2_/L in the bench-scale reactor). Average influent and effluent composition in the full-scale nitrifying activated sludge reactor is shown in Table 1.

### 2.3 Analytical methods

Total and soluble chemical oxygen demand (COD), total suspended solids (TSS), volatile suspended solids (VSS), alkalinity, total and soluble Kjeldahl nitrogen (TKN, sTKN), NH_4_^+^-N, combined NO_3_+NO_2_-N (NO_X_-N), total phosphorus, and orthophosphate were monitored 3 to 5 times/week in influent and effluent samples, per Standard Methods (APHA, 2005).

### 2.4 Biomass sampling and DNA extraction

Reactor biomass was archived weekly to biweekly for PCR and sequencing-based analyses. Six 1 mL aliquots of mixed liquor were centrifuged at 10,000g for 3 minutes, and the supernatant was decanted and replaced with 1 mL of tris-EDTA buffer. The biomass pellet was then resuspended, and the aliquots were centrifuged again at 10,000g for 3 minutes and the supernatant decanted, leaving only the biomass pellet to be transferred to the −80°C freezer. All samples were kept at −80°C until DNA extraction was performed with the FastDNA SPIN Kit for Soil (MPBio, Santa Ana, CA, USA) per the manufacturer’s instructions.

### 2.5 Quantitative PCR and comammox *amoA* cloning

Quantitative PCR (qPCR) assays were performed targeting AOB *amoA* via the *amoA-*1F and *amoA*-2R primer set (Rotthauwe et al., 1997), Nitrospira *nxrB* via the 169f /638R primer set (Pester et al., 2014), comammox *amoA* via the Ntsp-amoA 162F/359R primer set (Fowler et al., 2018), AOA *amoA* via the Arch-amoAF/AR primer set (Francis et al., 2005), and total bacterial (universal) 16S rRNA genes via the Eub519/Univ907 primer set (Burgmann et al., 2011). All assays employed thermocycling conditions reported in the reference papers, and were performed on a Bio-Rad C1000 CFX96 Real-Time PCR system (Bio-Rad, Hercules, CA, USA). Details on target genes, reaction volumes and reagents can be found in Supporting Information. After each qPCR assay, the specificity of the amplification was tested with melt curve analysis and agarose gel electrophoresis.

Comammox *amoA* genes were amplified, cloned, and sequenced on day 407 to generate standards for qPCR and confirm specificity of the comammox primer set, following previously described methods (Wells et al., 2009) (See Supporting Information for details).

### 2.6 16S rRNA, *amoA,* and *nxrB* gene amplicon sequencing

16S rRNA and functional gene amplicon library preparations were performed using a two-step multiplex PCR protocol, as previously described (Griffin and Wells, 2017). All PCR reactions were performed using a Biorad T-100 Thermocycler (Bio-Rad, Hercules, CA). The V4-V5 region of the universal 16S rRNA gene was amplified in duplicate from 27 dates collected over the course of reactor operation using the 515F-Y/926R primer set (Parada et al., 2016). To characterize the overall *Nitrospira* and comammox *Nitrospira* microdiversity in the system, *Nitrospira nxrB* gene amplicons were sequenced from duplicate biological samples from day 407 and comammox *amoA* gene amplicons were sequenced from duplicate biological samples of 6 time points (days 229, 262, 291, 370, 383, and 407) using primers mentioned in section 2.5 (Fowler et al., 2018; Pester et al., 2014). Further details on thermocycling conditions and primer sequences can be found in Supporting Information.

All amplicons were sequenced using a MiSeq system (Illumina, San Diego, CA, USA) with Illumina V2 (2×250 paired end) chemistry at the University of Illinois at Chicago DNA Services Facility and deposited in GenBank (accession number for raw data: PRJNA480047; also see Table S2). Procedures for sequence analysis and phylogenetic inference can be found in Supporting Information.

### 2.7 Quantitative Fluorescence *in situ* Hybridization (qFISH)

Reactor biomass was fixed and archived biweekly to monthly for probe-based qFISH analyses, following previously described methods (Wells et al., 2017). Briefly, reactor biomass samples were fixed in a 4% formaldehyde solution for two hours at 4°C followed by storage at −20°C in a 1:1 solution of phosphate buffered saline (PBS) and ethanol. qFISH was performed to estimate the relative abundance of canonical *Nitrospira*, canonical ammonia oxidizing bacteria (AOB), putative comammox *Nitrospira*, and total bacteria (see Supporting Information and Table S1 for probe sequences, staining procedure, imaging, and quantification methods).

## 3 Results

### 3.1 Bioreactor performance

The bench-scale reactor was initially assessed for its ability to remove total inorganic nitrogen (TIN) via the partial nitritation/anammox (PN/A) pathway. Effluent N concentrations over time are shown in Figure 1. The greatest TIN removal performance was observed in the first 77 days of operation after inoculation with sidestream PN/A biomass, where 49 ± 12% of TIN and 78 ± 17% of NH_4_^+^ was removed from the reactor. A relatively low average ratio of NO_3_^−^ production to NH_4_^+^ removal of 0.24 gNO_3_^−^-N/gNH_4_^+^-N during this time indicated moderate NOB suppression, as the theoretical nitritation-anammox pathway with no NOB activity produces a ratio of 0.11 gNO_3_^−^- N/gNH_4_^+^-N. However, the activity and biomass of slow-growing anammox were rapidly lost from the system. Batch activity testing revealed that, by day 34, the maximum potential anammox activity had declined by 88% (Figure 2), and never appreciably recovered over the course of >400 days of operation. Subsequent molecular profiling demonstrated a parallel steep decline in anammox abundance (described below).

**Figure 1.**
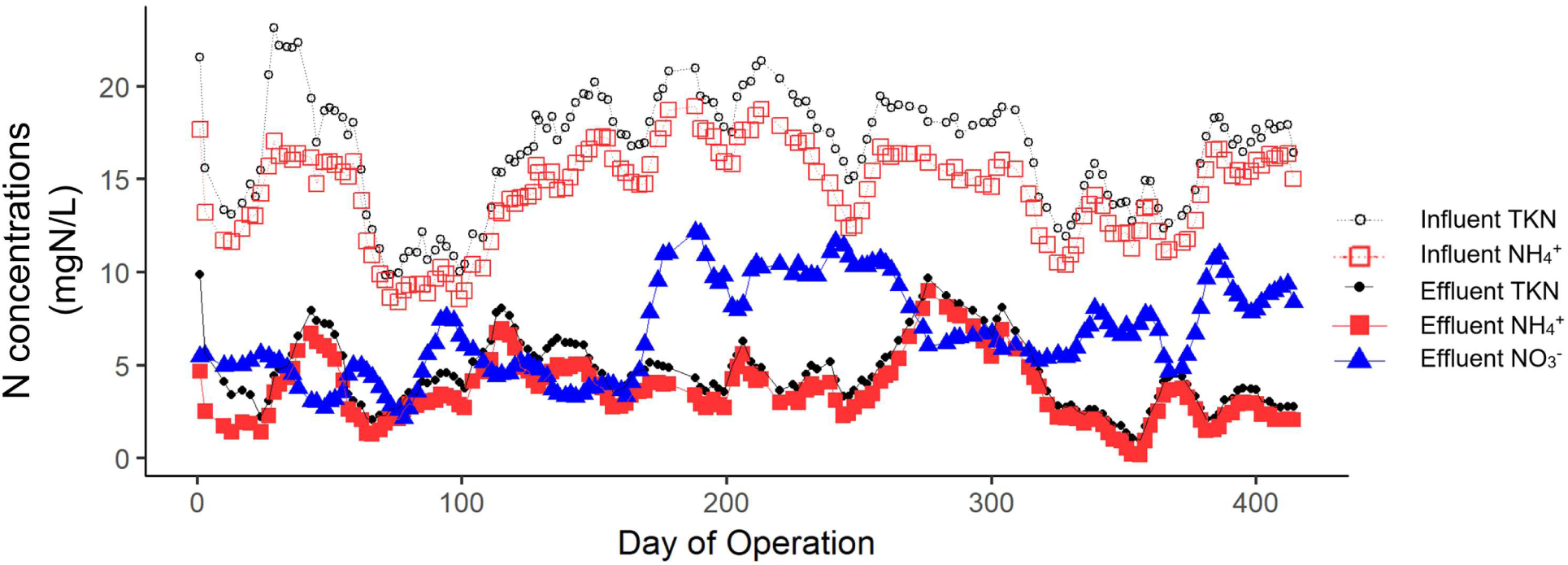
Nitrogen concentrations in reactor influent and effluent over time. All parameters are shown as a two-week rolling average. “NO_3_^−^” was measured as NO_3_^−^ + NO_2_^−^, but weekly effluent NO_2_^−^ measurements revealed that NO_2_^−^ comprised <1% of NO_3_^−^ + NO_2_^−^ on average (max 6%).

**Figure 2.**
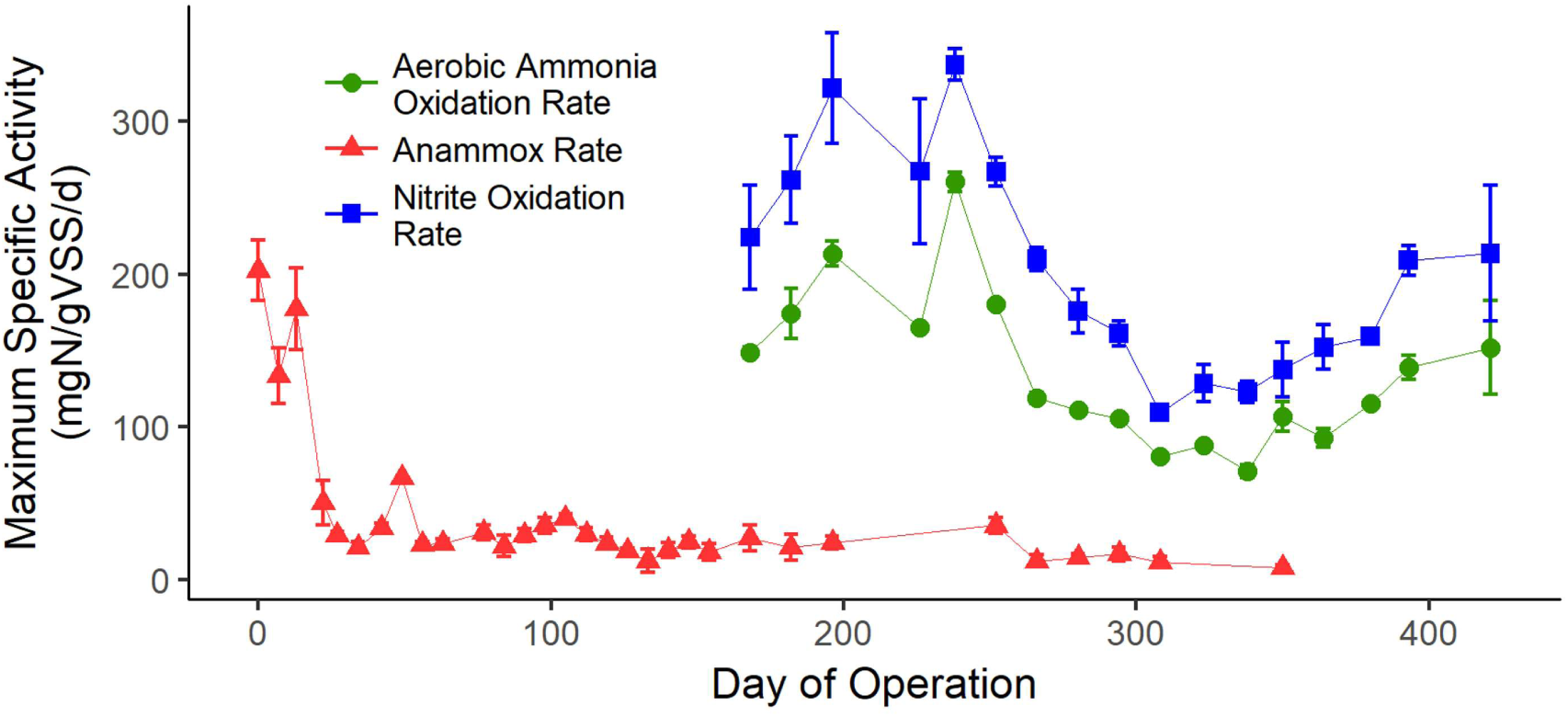
Maximum specific activities for aerobic ammonia oxidation rate, nitrite oxidation rate, and anaerobic ammonia oxidation (anammox) rate in reactor biomass over time from batch assays. Rates were measured under non-limiting substrate concentrations.

Loss of anammox activity and abundance coincided with a decrease in TIN removal, but high rate complete NH_4_^+^ oxidation was maintained in the presence of low DO with intermittent aeration. Between day 77 and the end of the study (day 414), TIN removal was 21 ± 19%, less than half of that of the initial 77 days. During the same time period (day 78-414), NH_4_^+^ removal did not decrease significantly, and averaged 74 ± 17%. The length of intermittent aeration intervals was adjusted with changes in influent strength during this time period, and varied between 1 to 2 minutes of aeration in 2 to 30-minute intervals. Target peak DO was between 0.2 – 1.0 mgO_2_/L. This aeration strategy was originally chosen as a hypothesized means to suppress NOB activity and therefore achieve (partial) nitritation (Gilbert et al., 2014). However, complete nitrification of NH_4_^+^ to NO_3_^−^, rather than nitritation of NH_4_^+^ to NO_2_^−^, was consistently achieved from days 78-414. NO_2_^−^ accumulation in the effluent was not observed throughout the study, with average effluent NO_2_^−^ of 0.07 ± 0.04 mgNO_2_^−^-N/L (less than 1% of total effluent NO_2_^−^ + NO_3_^−^). Maximum aerobic NH_4_^+^ oxidation and NO_2_^−^ oxidation rates were first measured on day 168, and revealed a greater NO_2_^−^ oxidation rate (Figure 2). This was consistently maintained throughout the experimental period, with an average ratio of 1.5:1 maximum NO_2_^−^ oxidation to NH_4_^+^ oxidation rate.

The bench scale low DO nitrifying SBR was operated in parallel to a full-scale conventional high DO nitrifying activated sludge bioreactor at the O’Brien WRP, offering an opportunity to compare performance characteristics between these systems. The O’Brien WRP achieved lower average effluent NH_4_^+^ values of 0.7 ± 0.1 mgNH_4_^+^-N/L compared to 3.6 ± 2.6 mgNH_4_^+^-N/L for the bench-scale low DO reactor (Table 1), in large part because a high effluent NH_4_^+^ residual was intentionally selected for the bench-scale system as a putative NOB out-selection strategy (Regmi et al., 2014) for PN/A operation. When the bench-scale reactor HRT was fixed at 9 hours from days 0 – 366, the NH_4_^+^ loading and removal rate was greater in the full-scale system (∼42 and 40 mgNH_4_^+^-N/L/d, respectively) than in the bench scale system (40.1 and 28.8 mgNH_4_^+^-N/L/d, respectively). However, once PLC control of HRT was implemented on day 358, the average NH_4_^+^ removal rate of 58.6 mgNH_4_^+^-N/L/d in the bench-scale reactor exceeded that of the full-scale high DO system (∼38 mgNH_4_^+^-N/L/d) from days 358 – 414. This was despite operation at significantly lower DO; the bench-scale reactor utilized intermittent aeration with a peak DO of ∼1 mgO_2_/L (effectively ranging between 0 and 1 mgO_2_/L throughout the react period), while the full-scale system utilized constant aeration with DO ranging from 0 mgO_2_/L at the beginning to 3 – 5 mgO_2_/L at the end of the aeration basins. The temperature of the bench-scale reactor during days 358 – 414 was only slightly higher than the full-scale system: 20.8 ± 0.8°C vs. 18.8 ± 1.5°C, respectively.

### 3.2 16S rRNA sequencing demonstrates strong enrichment of *Nitrospira* and decline in AOB

Results from 16S rRNA amplicon gene sequencing analyses over the course of reactor operation demonstrated a decline of anammox (which included all OTUs affiliated with the genus *Brocadia*; no other anammox genera were detected) relative abundance from 4% of total amplicons on day 13 to less than 0.1% by day 161 (Figure 3). This decline paralleled the rapid decline in maximum anammox activity (Figure 2) and TIN removal (Figure 1) in this system. While NOB abundance was expected to exceed that of AOM based on the maximum activity ratio (Figure 2) and the lack of a NO_2_^−^ residual, 16S rRNA sequencing revealed an unexpected strong declining trend of canonical betaproteobacterial AOB (OTUs affiliated with the genus *Nitrosomonas*; no *Nitrosospira*, AOA, or gammaproteobacterial AOB were detected) from 4.4% of total reads on day 3 to 0.8% of total reads by day 407, while putative NOB (which included only OTUs affiliated with the genus *Nitrospira*; *Nitrobacter* and *Nitrotoga* were not detected, and *Nitrolancea* was <0.2% for all time points) steadily climbed from 3.1% of total reads on day 3 to an unusually high 53% of reads on day 407 (Figure 3). By clustering the *Nitrospira*-affiliated 16S rRNA gene sequences at the identity cutoff of 97%, the micro-diversity of the *Nitrospira* in the community appeared to be very low: just two OTUs (OTU2 and OTU4) contributed 95.4 – 99.8% (depending on time point) of all *Nitrospira* 16S rRNA gene amplicons. Around day 100 the less abundant of these two, OTU2, began to accumulate (Figure 3) and by day 407 accounted for 92% of total NOB and 49% of total 16S rRNA reads. This trend coincided with the strong decrease in canonical AOB abundance (Figure 3).

**Figure 3.**
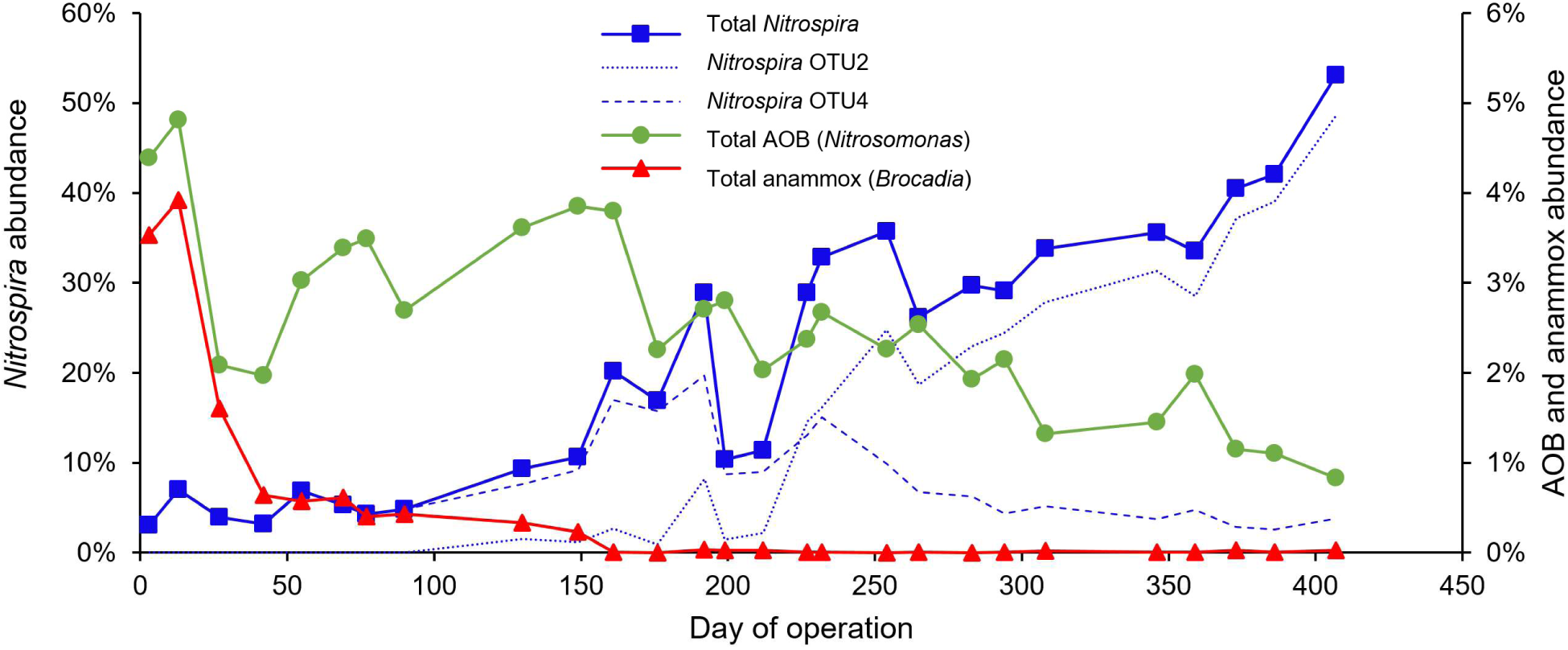
Relative abundance of total *Nitrospira*, total AOB (all *Nitrosomonas*) and total anammox (all *Brocadia*), based on 16S rRNA amplicon sequence analysis. Relative abundance for *Nitrospira* is shown on the left y-axis, and AOB and anammox are shown on the right y-axis. *Nitrospira* OTU2 and OTU4 together comprise 95.3 – 99.8% of total *Nitrospira* for all time points.

Taken together, these patterns of relative abundance of *Nitrospira* and AOB-affiliated OTUs in 16S rRNA gene sequencing datasets led us to hypothesize the accumulation of comammox *Nitrospira* due to the consistent high ammonia oxidation activity and complete nitrification in the reactor. Moreover, for the period that maximum aerobic ammonia oxidation rates were measured (days 168 – 414), the average ratio of relative abundance of *Nitrospira* to canonical AOB was 19:1 according to 16S rRNA amplicon sequencing results, much higher than the aforementioned 1.5:1 maximum nitrite oxidation: ammonia oxidation rate. This finding reaffirmed the hypothesis that comammox may play a dominant role in ammonia oxidation in this low DO nitrification reactor. This hypothesis was further reinforced by a closer investigation of the phylogenetic affiliation of the amplicon sequence variants (ASV) in *Nitrospira* OTU2 and OTU4 (Figure S3), which suggested that ASV within OTU2 were phylogenetically affiliated with the 16S rRNA gene of *Ca.* Nitrospira nitrosa.

### 3.3 Functional gene quantification reveals dominance of comammox over canonical AOB

The presence or abundance of comammox *Nitrospira* cannot be resolved from 16S rRNA amplicon sequencing alone (Lawson and Lücker, 2018), and the unusually abundant *Nitrospira* population warranted the use of better tools for functional group quantification. For this purpose, qPCR was used to quantify comammox (targeting *amo*A), canonical AOB (*amo*A), total *Nitrospira* (*nxrB*), and total bacteria (16S rRNA). Ammonia oxidizing archaea were also investigated via endpoint and qPCR targeting archaeal *amoA* genes, but were not detected on any samples dates based on the lack of detection or target bands in gel electrophoresis of qPCR products. *Nitrobacter* and *Nitrotoga* NOB were absent from 16S rRNA sequencing analysis and thus were not investigated with qPCR.

qPCR measurements confirmed broad NOB and AOB trends observed in the 16S rRNA sequencing results (Figure 4). Based on qPCR results, the average ratio of total *Nitrospira* to AOB for days 168 – 414 was 66:1 (Nitrospira *nxrB*: AOB *amoA*), higher than the 19:1 ratio from 16S rRNA sequencing and the 1.5:1 ratio from maximum activity tests. The relative abundance of total *Nitrospira* based on qPCR measurements (*Nitrospira nxrB* normalized to total bacterial 16S rRNA genes) increased over time to 55% on day 407, confirming trends and the surprisingly high total *Nitrospira* relative abundance we observed in 16S rRNA sequencing datasets at this time point (53%).

**Figure 4.**
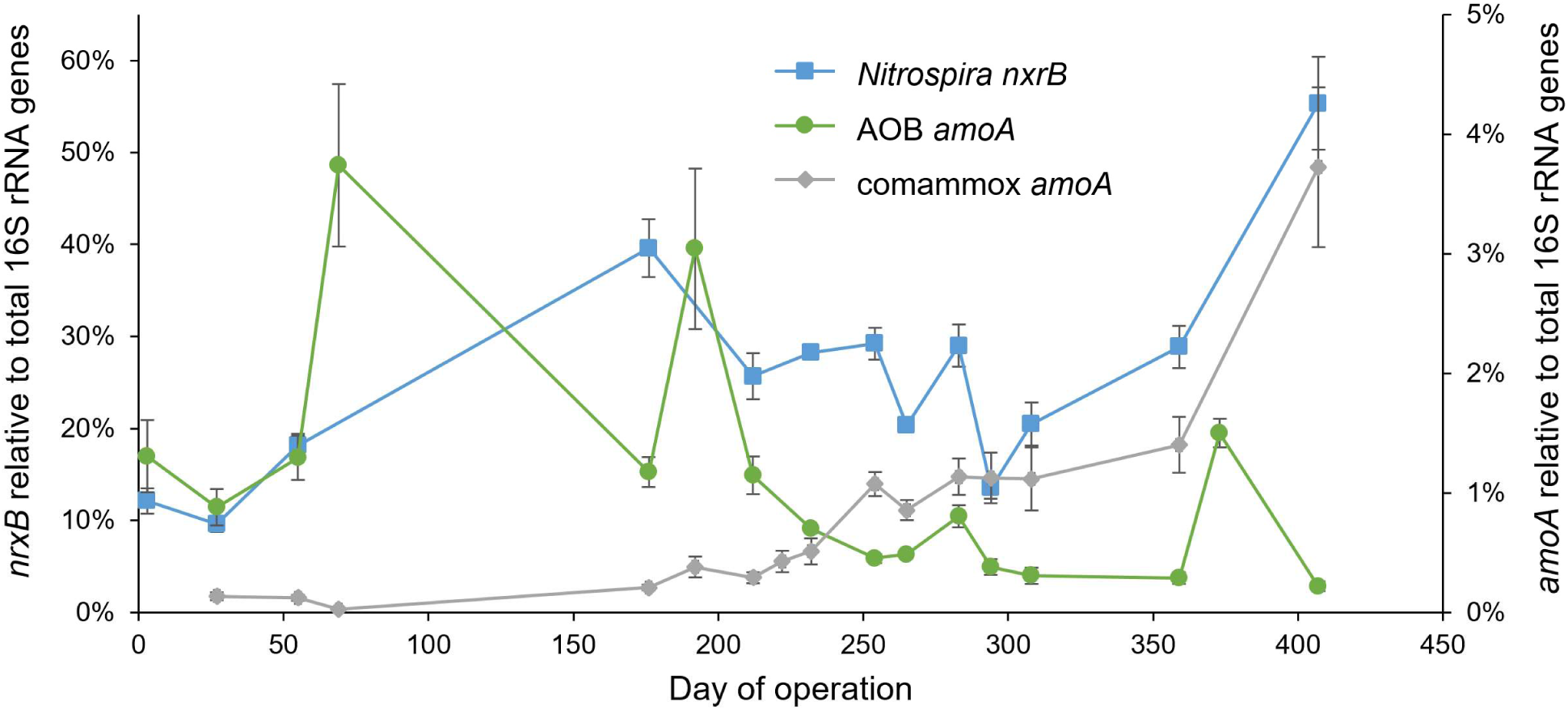
Abundance of *Nitrospira nxrB* (squares), AOB *amoA* (circles), and comammox *amoA* (diamonds), as measured by qPCR. Results are normalized to total bacterial 16S rRNA gene copies. *Nitrospira* results are shown on the left y-axis, and AOB and comammox results are shown on the right y-axis.

Based on qPCR measurements, comammox was the dominant ammonia oxidizer from days 254 – 414 of reactor operation, with an average 79% of total *amoA*. By day 407, comammox comprised 94% of all detected *amoA*. qPCR measurements targeting comammox *amoA* revealed an increasing abundance in comammox *Nitrospira* beginning around day 222 (Figure 4), and a concurrent decline in AOB *amoA* abundance.

This increasing abundance of comammox *Nitrospira* coincided closely with the increase in the *Nitrospira* OTU2 from 16S rRNA sequencing (Figure 3), though the relative abundances did not match, suggesting that *Nitrospira* microdiversity was more complex than the two dominant *Nitrospira* OTUs (97% identity) documented via analysis of 16S rRNA sequence data. While *Nitrospira* OTU2 from 16S rRNA sequencing data accumulated to 49% of total reads by day 407, comammox *amoA* accumulated to just 4% of total bacteria when measured relative to total bacterial 16S rRNA genes via qPCR. Interestingly, between days 254 and 414 of reactor operation, comammox *amoA* reads averaged just 5% relative to total *nxrB* reads. This result suggests that while comammox *Nitrospira* were the putative dominant ammonia oxidizers in this system (based on abundance relative to canonical AOB), a large fraction of the total *Nitrospira* community was incapable of comammox metabolism.

To verify specificity of comammox *amoA* primers and generate associated qPCR standards, three clones were generated from comammox *amoA* PCR products, sequenced, and phylogenetically analyzed. This analysis was employed for an initial assessment of relationship of comammox in this system to known comammox taxa. The results confirmed that all three clones affiliated with comammox *amoA* genes of comammox clade A and were most closely related to *Candidatus* N. nitrosa (van Kessel et al., 2015) and *Nitrospira* sp. UW LDO 01 (Camejo et al., 2017) (Figure 5b).

**Figure 5.**
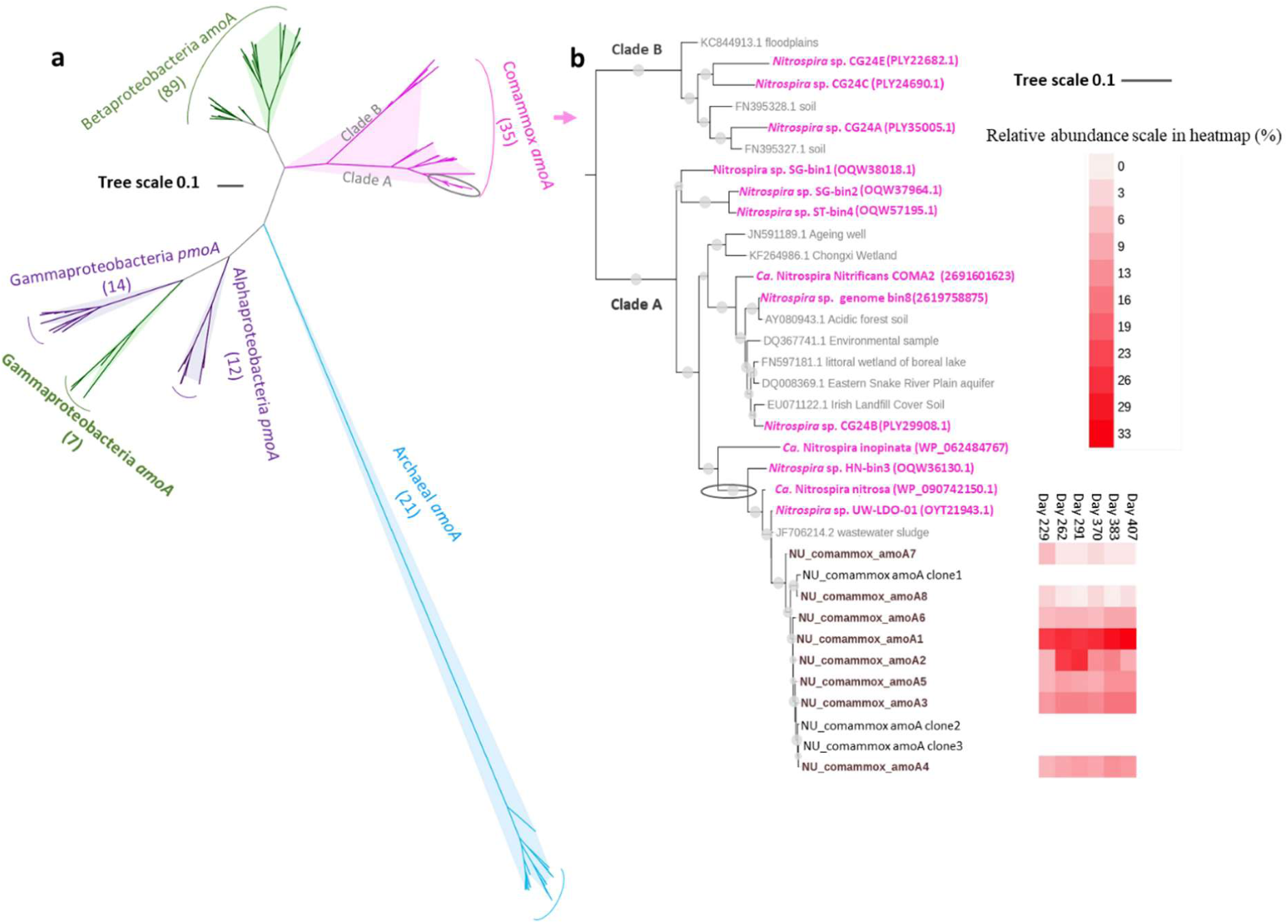
Phylogenetic affiliation of comammox *amoA* gene sequences from the low DO nitrification reactor via cloning and sanger sequencing and via high throughput amplicon sequencing (173 bp fragment). Scale bars indicate estimated change per nucleotide sequence. **a**) Maximum likelihood tree showing the phylogenetic relationship of the 3 clone sequences and 8 amplicon sequence variants (ASV) from sequencing (circled in above tree) with other *amoA* gene superfamily members (167 sequences). **b**) Maximum likelihood tree showing phylogenetic relationship between the 3 comammox *amoA* clone sequences and 8 comammox *amoA* ASV from amplicon sequencing (black, names beginning with NU_comammox) with NCBI database-derived comammox *amoA* gene sequences. The *amoA* gene from *Nitrosococcus watsonii* C-113 was applied as the outgroup. The heatmap indicates the relative abundance of the 8 comammox *amoA* ASV we detected in reactor biomass at the six time-points. Genome-derived comammox *amoA* genes are in magenta and other database-derived comammox *amoA* genes are in light gray. Nodes with the bootstrap value > 70% have all been marked with a gray circle, and the size of the circle is in positive correlation with the bootstrap value.

### 3.4 qFISH confirms *Nitrospira* enrichment

qFISH measurements of reactor biomass from three sample dates was used as a check on the trends and functional group abundances as observed by qPCR. The trend of the increasing ratio of NOB:AOB was confirmed; this ratio increased from 0.8:1 on day 35 to 15:1 on day 398 (Figure 6). A composite ratio reveals an enrichment of NOB from < 1% on day 35 to 35% of the overall bacterial community on day 398. This result confirms the strong increasing trend in total NOB documented via both 16S rRNA sequencing and qPCR analyses.

**Figure 6.**
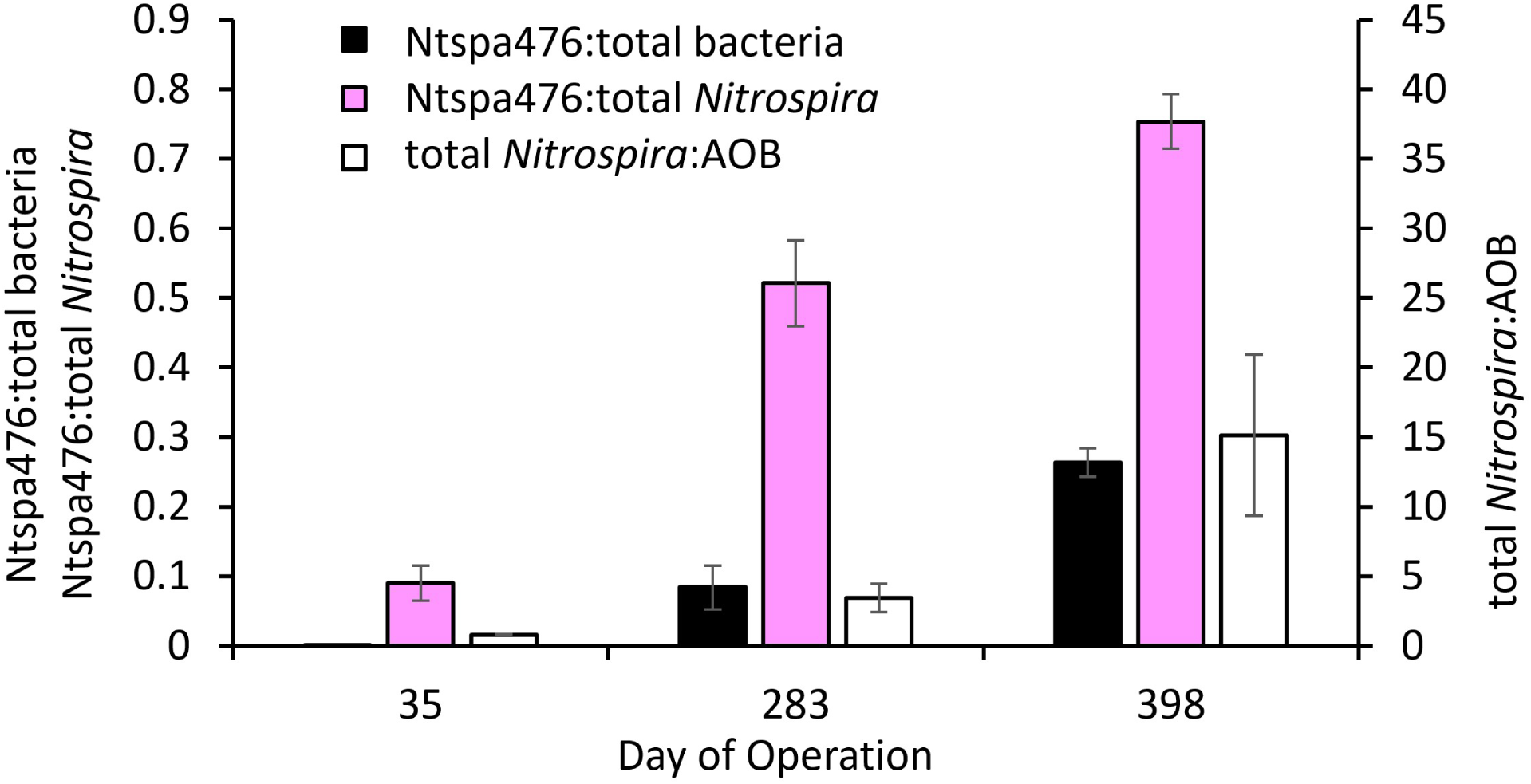
qFISH results of Ntspa476 probe (see text section **3.4** and van Kessel et al. [2015a]), total Nitrospira, total AOB, and total bacteria. See Supporting Information regarding the specificity of the Ntspa476 FISH probe to comammox.

The FISH probe Ntspa476 utilized in this study was previously developed to target putative comammox *Ca.* N. nitrosa and *Ca.* N. Nitrificans (van Kessel et al., 2015a). The specificity and coverage of this probe set to our specific system was evaluated before conducting the qFISH assays (see Supporting Information). In brief, our analysis suggests that the probe is not capable of reliably resolving comammox from conventional nitrite-oxidizing *Nitrospira*; the inability of 16S rRNA gene-based methods to do so has been raised before (Lawson and Lücker, 2018). However, our analysis suggests that Ntspa476 primarily targets lineage II *Nitrospira* that include comammox *Nitrospira*, and so provide a finer resolution method for confirming trends in this lineage. 75% of the total *Nitrospira* on day 398 hybridized with Ntspa476 (Figure 6), substantially higher than the results of the qPCR analysis for comammox, which show 7% on day 407 as a fraction of total *nxrB* (Figure 4). These results and the analysis in Supporting Information suggest that a portion of *Nitrospira* targeted by Ntspa476 and prevalent in this reactor are non-comammox *Nitrospira*. The function of the surprisingly high abundance and putatively non-comammox *Nitrospira* taxa in this reactor warrants further investigation.

### 3.5 Functional gene sequencing

As described above, 16S rRNA sequencing suggested that a single OTU dominated the *Nitrospira* community by day 407 (Figure 3), while qPCR demonstrated that comammox comprised 7% of the *Nitrospira* community at this time point (Figure 4). To better resolve microdiversity within the *Nitrospira* community, *nxrB* functional gene sequencing was performed on the sample from day 407. 22,440 total *nxrB* sequences were generated from which 30 amplicon sequence variants (ASV) were identified with DADA2 (Callahan et al., 2016). The resulting *nxrB* phylogenetic tree (Figure 7) confirms that the *nxrB* gene cannot be reliably applied to distinguish comammox from canonical *Nitrospira* (i.e. see the top branch of Figure 7 for interspersed comammox and canonical *Nitrospira;* also see Lawson and Lücker [2018]), but still offers insight into the diversity of the *Nitrospira* community. 15% of the Nitrospira *nxrB* sequences clustered with the *nxrB* gene in the comammox genomes of *Ca.* N. nitrosa and *Nitrospira* sp. UW LDO 01. Another 37% of *nxrB* sequences clustered with genomes of the canonical NOBs *Nitrospira defluvii* and *Nitrospira* sp. Strain ND1, of which two strains have been reported with a microaerophilic or hypoxic ecological niche (Lucker et al., 2010; Ushiki et al., 2017). Two additional clusters of *nxrB* sequences accounting for 10% and 38% of the total *nxrB* amplicons do not affiliate with currently available genomes, but a blast search in NCBI returned three sequences (indicated with “[*accession #*] activated sludge” labels in Figure 7) with 92 – 98% identity to the *nxrB* genes in these two clusters.

**Figure 7.**
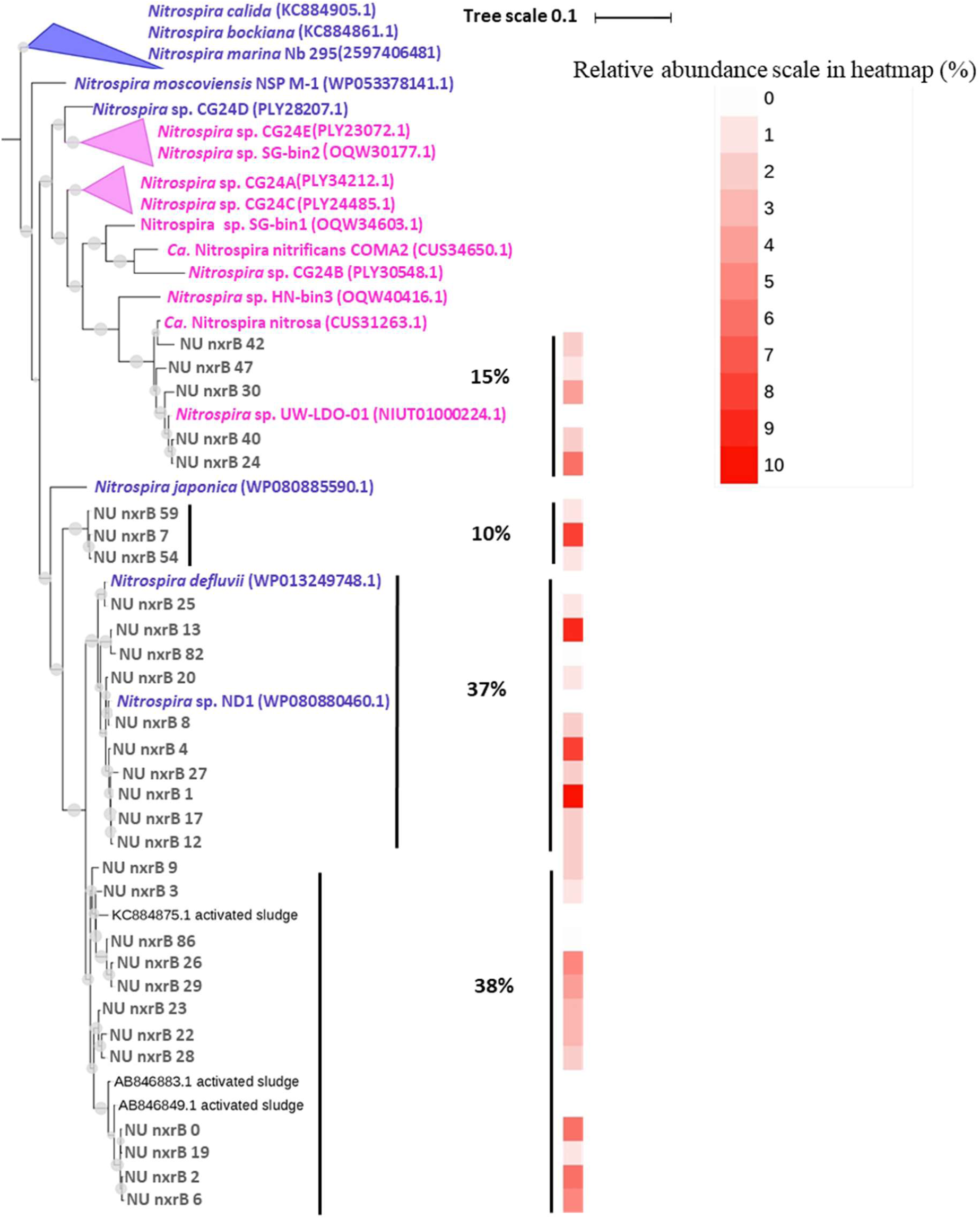
Maximum-likelihood phylogenetic tree of *Nitrospira nxrB* gene sequences (453 bp fragment). The *nxrB* gene from *Nitrobacter hamburgensis* X14 was applied as the outgroup. Nodes with the bootstrap value > 70% have all been marked with a gray circle, and the size of the circle is in positive correlation with the bootstrap value. 30 *nxrB* ASV recovered in this study on day 407 of reactor operation are shown in bold as “**NU nxrB ##**”. In magenta are *nxrB* genes from known comammox *Nitrospira* genomes, and in purple are *nxrB* genes of known non-comammox *Nitrospira* genomes. Percentages to right of the tree indicate relative abundance of *nxrB* clusters relative to total *nxrB* in the low DO nitrification reactor. The red to white heatmap indicates the percentage of each **NU nxrB** ASV relative to the overall *Nitrospira* in the reactor. No *nxrB* genes from currently available genomes cluster with the 10 and 38% abundance **NU nxrB** clusters, so the three closest non-genome database sequences are shown as “[*accession #*] activated sludge”.

Functional gene sequencing of the *amoA* gene of comammox *Nitrospira* on 6 sampling points from days 229 – 407 generated between 28,194 and 57,637 (depending on the time point) total *amoA* sequences from which 217 ASV were identified, and 8 of them accounted for 94 – 99% of the total. Phylogenetic analysis revealed that these 8 sequences cluster within comammox Clade A (see “NU_comammox_amoA” samples in Figure 5b). The close association of the comammox population in the low DO nitrification SBR with *Nitrospira* sp. UW LDO 01 (Camejo et al., 2017) and *Ca.* Nitrospira nitrosa (van Kessel et al., 2015a) suggested by *nxrB* sequencing (Figure 7) and sequencing of clones generated as qPCR standards (Figure 5b) was confirmed by this high throughput comammox *amoA* sequence analysis.

## 4. Discussion

In this study, comammox *Nitrospira* were observed to accumulate over time in a low DO nitrification reactor treating mainstream municipal wastewater to eventually become the numerically dominant ammonia oxidizer as confirmed by sequencing, qPCR, and microscopy-based methods. This counters the prevailing assumption that comammox play a minor role in wastewater treatment bioreactors (Annavajhala et al., 2018; Gonzalez-Martinez et al., 2016). At least two previous studies have challenged this same assumption. In the original publication of the discovery of *N. inopinata* (Daims et al., 2015), metagenomic analysis revealed that comammox comprised 43 to 71% of the total *Nitrospira* population at the WWTP of the University of Veterinary Medicine in Vienna, Austria. A subsequent effort to develop comammox-specific *amoA* qPCR primers revealed comammox at the same plant as comprising 14 to 35% of total *amoA* via qPCR (Pjevac et al., 2017), still a minority of the overall NH_4_^+^ oxidizing community. In a separate study, while developing a bench-scale biological nutrient removal reactor with synthetic feed, researchers at the University of Wisconsin-Madison serendipitously enriched comammox to 38% of total *Nitrospira*, or 5.4% of total normalized metagenomic reads during the first stage of operation (Camejo et al., 2017). Comammox were far more abundant than AOA and AOB in this stage (which together comprised just 0.23% of total reads), suggesting that comammox was the dominant NH_4_^+^ oxidizer. The enrichment was transient, however, as subsequent sampling revealed its absence, and was associated with synthetic rather than real wastewater feed.

Significantly, this study is the first to show comammox as the dominant NH_4_^+^ oxidizer in a reactor using actual wastewater as influent, demonstrating that comammox may play an important role in biological nutrient removal systems in practice under certain conditions. Comammox *Nitrospira* was not detected in the parallel O’Brien WRP full-scale high DO nitrifying activated sludge system, indicating that operating parameters such as low DO and high SRT may be important for their selection. Of the two studies that have found comammox to (at least transiently) dominate the ammonia oxidizing community of a biological nutrient removal reactor – the present study and Camejo et al., 2017, both of which selected for closely related comammox strains within Clade A – a few key similarities between the two SBRs stand out:

- Low in-reactor N concentrations: 0 –12 mgN/L of NH_4_^+^, NO_3_^−^, and NO_2_^−^(Camejo et al., 2017), and 0 – 14 mgN/L of NH_4_^+^ and NO_3_^−^, 0 – 0.2 mgN/L of NO_2_^−^ (present study)
- Low DO: 0 – 0.4 mgO_2_/L with constant aeration (Camejo et al., 2017), and 0 – 1.0 mgO_2_/L with intermittent aeration (present study)
- Very high average SRT: 80 days (Camejo et al., 2017) and 99 days (present study)

The above observations are potentially unfavorable for applications targeting shortcut N removal processes, including PN/A. Low DO and high SRT are required to retain anammox activity and biomass in PN/A systems, but our results suggest that these conditions may inadvertently select for comammox when applied to the relatively low N concentrations typical of mainstream wastewater. However, while any NO_2_^−^ oxidation is considered unfavorable in PN/A systems, it is possible that the presence of comammox may still be compatible with their operation. Transient NO_2_^−^ accumulation produced by *N. inopinata* during oxidation of NH_4_^+^ has been observed (Daims et al., 2015; Kits et al., 2017), and FISH imaging revealed comammox *Candidatus* N. nitrificans and *Candidatus* N. nitrosa in co-aggregation with *Brocadia*-affiliated anammox (and in the absence of canonical AOM) (van Kessel et al., 2015a). While this implies that comammox may be compatible with well-functioning PN/A systems, such systems may require a more careful control of redox conditions to ensure that reduction of NO_2_^−^ by anammox is favored over NO_2_^−^ oxidation by comammox or canonical NOB. Further studies involving the coordinated activity and cross-feeding of comammox and anammox will be required to better delineate these conditions.

In contrast, there are no obvious disadvantages to the presence of comammox in low DO nitrification systems, as in the present study. In fact, comammox may be especially suited to systems targeting very low effluent NH_4_^+^ levels due to their high substrate affinity (Kits et al., 2017). In the present study, the NH_4_^+^ oxidation rate in the bench-scale reactor exceeded that of the full-scale system when variable HRT was utilized on days 358 – 414, despite operation at much lower DO. This suggests that comammox *Nitrospira* may be well-suited to energy-efficient methods for complete ammonia oxidation, and thus offer an alternative to high-DO conventional nitrification systems. Low DO systems for nitrification have been shown to save up to 25% in energy use over conventional, high-DO systems (Keene et al., 2017) without sacrificing process stability (Fitzgerald et al., 2015; Park and Noguera, 2007, 2004).

### 4.1 High NOB to AOM ratios

A discrepancy between nitrite and ammonia oxidation rates and nitrite and ammonia oxidizing organism abundance in nitrifying activated sludge biomass, as in this study, has been observed before (Fitzgerald et al., 2015; Schramm et al., 1999; Wang et al., 2015). Wang et al. observed NOB:AOB ratios of anywhere from 9:1 to 5000:1 in five activated sludge reactors as measured by qPCR, and Schramm et al. observed an average NOB:AOB ratio of 24:1 on the surface of biomass aggregates as measured by FISH. *Nitrospira* was the most abundant NOB in both studies. Other researchers have speculated that undetected comammox *Nitrospira* may fill this gap (Daims et al., 2015), but even in this study, their presence does not fully explain the difference as measured by qPCR, as non-comammox *Nitrospira* still comprised ∼50% of total bacteria by day 407, with an NOB: AOM ratio of about 14:1 on day 407 if comammox *Nitrospira* are counted as both NOB and AOM.

The most common explanations for high NOB:AOM ratios are (1) oversimplification of the metabolic versatility of NOB, particularly for *Nitrospira*, and (2) under-estimation of AOM. With regards to explanation (1), the *Nitrospira* genus displays an impressive diversity in terms of metabolic capabilities (Daims et al., 2016; Koch et al., 2015). Koch et al. showed that *Nitrospira moscoviensis* contains genes encoding for urease and formate dehydrogenase, the latter of which need not be tied to nitrite oxidation. Daims et al., in their review of *Nitrospira* metabolic versatility, pointed out that *Nitrospira* are also capable of oxidizing hydrogen gas under oxic conditions and reducing NO_3_^−^ in the presence of simple organics under anoxic conditions. Given this versatility, a portion of *Nitrospira* may not be oxidizing NO_2_^−^ as their primary energy source. Additionally, NOB may proliferate in the presence of a nitrite oxidation/nitrate reduction loop in conjunction with heterotrophs (Winkler et al., 2012), facilitated by oxic-anoxic zones in space or time, as with intermittent aeration in the present study.

Fitzgerald et al. (2015) suggest scenarios for the latter explanation (2), that AOM abundance may be underestimated. In a study of low DO nitrification systems, one nitrification reactor contained no known AOM despite the presence of complete nitrification to NO_3_^−^. Through batch experiments on pure culture isolates from the reactor they demonstrated NH_4_^+^ removal beyond typical assimilation for five organisms previously not known to oxidize NH_4_^+^. The researchers suggest that heterotrophic nitrification may be especially important for low DO systems. It should also be noted that it is possible that currently available comammox *amoA* primers may underestimate comammox abundance.

Understanding of the relevance of comammox to diverse BNR systems is clearly in early stages, and diverse research questions remain. Looking forward, a key need for nutrient removal researchers is for better specificity of maximum activity tests as performed in this study. Aerobic ammonia and nitrite oxidation as measured give a reasonable estimate of total potential oxidation rates, but do not distinguish between the activity of canonical AOM and comammox *Nitrospira*. Better delineation of *in-situ* activities of key functional groups is needed to characterize these systems, and could begin with measurements of gene expression within cycles.

## 5. Conclusions

- Comammox *Nitrospira* dominated the ammonia oxidizing community in a mainstream nitrification reactor fed with real municipal wastewater for >400 days. By the end of reactor operation, comammox *Nitrospira* accounted for 94% of the AOM community. This counters the notion that comammox are not relevant to wastewater treatment technologies.
- Efficient nitrification was demonstrated at low DO concentrations of 0.2 – 1.0 mg/L via intermittent aeration. Volumetric ammonium removal rates averaged 58.6 mgN/L-d during the final two months of operation when comammox abundance was particularly high. These rates were higher than in a parallel full-scale high DO (3-5 mg/L) nitrifying conventional activated sludge reactor, suggesting the potential for an energy-efficient and comammox-driven low DO alternative to conventional high-DO nitrification processes.
- *Nitrospira* increased in abundance during reactor operation to 53% of the overall microbial community. The presence of comammox does not fully explain the observed very high abundance nor high ratios of *Nitrospira* to AOM. Further research is needed to investigate the metabolic versatility within the *Nitrospira* genus and functional importance to reactor operation.
- Operational conditions (low DO, low NH_4_^+^, and high SRT) in this reactor mirror those commonly used or undergoing testing in mainstream partial nitritation/anammox reactors, suggesting that efforts to cultivate shortcut N removal bioprocesses in the mainstream may inadvertently select for comammox. These results further indicate that comammox may play an increasingly important role in low DO nutrient removal biotechnologies.

## 6. Acknowledgements

Many thanks to Lachelle Brooks, Jianing Li, Qiteng Feng, Christian Landis, Adam Bartecki, and George Velez for help with reactor operation, sampling, and activity testing. We also thank MWRD staff and operators for site support at O’Brien WRP.

### Funding

This study was funded by the Metropolitan Water Reclamation District of Greater Chicago and the National Science Foundation Graduate Research Fellowship under Grant No. DGE-1324585.

