## Supporting Information for "Comammox *Nitrospira* are the dominant ammonia oxidizers in a mainstream low dissolved oxygen nitrification reactor"

### S1. Methods

#### S1.1 SRT calculation

No intentional wasting occurred throughout the study, so the average (± standard deviation) solids retention time (SRT) of 99 ± 44 days was determined by effluent solids losses and reactor mixed liquor sampling. Effluent solids losses were minimized by 26-L downstream clarifiers which were used to return effluent solids to the reactor. Samples for effluent solids measurements were therefore sampled from the overflow of these clarifiers. SRT as discussed in this manuscript was calculated with the following formula:

$$SRT = \frac{X_R V}{X_E Q_E + X_R Q_S} \qquad \textit{(Eq. 1)}$$

$SRT$ = Solids retention time

$X_R$ = Suspended solids concentration in the reactor

$V$ = Volume of reactor

$X_E$ = Suspended solids concentration in clarifier overflow

$Q_E$ = Effluent flow rate

$Q_S$ = Sampling "flow rate," i.e. volume & frequency of mixed liquor sampled from reactor

#### S1.2 Batch maximum activity assays

Batch kinetic assays were performed to determine maximum activities of anammox, ammonia-oxidizing microorganisms (AOM), and NOB functional groups under non-limiting substrate conditions, as previously described (Laureni et al., 2016). Anammox maximum activity tests were performed *in situ* at the end of reactor cycles when sCOD was at a minimum to avoid potential interference from denitrifiers. $NH_4^+$ and $NO_2^-$ were spiked to non-limiting conditions (approximately 15 mgN/L each) via ammonium chloride and sodium nitrite salt solutions and the reactor was mixed without aeration. Five to six grab samples were taken in 30-minute intervals and analyzed for $NH_4^+$, $NO_2^-$ and $NO_3^-$ by colorimetry (APHA, 2005).

According to stoichiometry from Strous et al.(Strous et al., 1998), the anammox metabolic pathway removes N (as nitrogen gas + biomass) at a ratio of 2.05 moles N per mole $NH_4^+$ removed:

$$1\ NH_4^+ + 1.32\ NO_2^- + 0.066\ HCO_3^- + 0.13\ H^+ \rightarrow$$
$$1.02\ N_2 + 0.26\ NO_3^- + 0.066\ CH_2O_{0.5}N_{0.15} + 2.03\ H_2O$$

Anammox activity as N removal (in mg N/L/d) was therefore calculated as 2.05 times the slope of the $NH_4^+$ drawdown curve. Only linear trends with $R^2$ values above 0.8 were used. Stoichiometric ratios of $NO_2^-$ drawdown and $NO_3^-$ production to $NH_4^+$ drawdown was checked against anammox stoichiometry to check that anammox was the dominant metabolic pathway in these batch assays.

AOM and NOB maximum activity assays were performed *ex situ* in duplicate 500-mL Erlenmeyer flasks, each containing 250 mL of mixed liquor. The flasks were placed on a shaker table with a water bath for temperature control between 19 – 21°C. DO was monitored with a Hach LDO® optical DO probe and were maintained at or above 3 $mgO_2$/L by shaking action and bubbling from small aquarium pumps. pH was monitored with the Hach PHC101® electrode and maintained between 7 and 8. $NH_4^+$ and $NO_2^-$ were spiked to non-limiting conditions (~20 $mgNH_4^+$-N/L and ~10 $mgNO_2^-$-N/L), and five grab samples were taken in 20-minute intervals and analyzed for $NH_4^+$, $NO_2^-$ and $NO_3^-$ by colorimetry (APHA, 2005).

For *ex situ* assays, AOM activity was taken as the slope of the $NH_4^+$ drawdown curve, and NOB was taken as the average of the (1) slope of the $NO_3^-$ production curve and (2) the sum of the slopes of $NH_4^+$ and $NO_2^-$ drawdown curves. Only linear trends with $R^2$ values above 0.8 were used.

### S1.3 qPCR gene targets, supermix and reaction conditions

qPCR gene targets included the AOB *amoA* gene (encoding the alpha subunit of ammonia monooxygenase in AOB) via the *amoA*-1F and *amoA*-2R primer set (Rotthauwe et al., 1997), the Nitrospira *nxrB* gene (encoding the beta subunit of nitrite oxidoreductase in *Nitrospira*) via the 169f /638R primer set (Pester et al., 2014), the comammox *amoA* gene (encoding the alpha subunit of ammonia monooxygenase specific to comammox *Nitrospira*) via the Ntsp-amoA 162F/359R primer set (Fowler et al., 2018), the AOA *amoA* gene (encoding the alpha subunit of ammonia monooxygenase specific to AOA) via the Arch-amoAF/AR primer set (Francis et al., 2005), and the total bacterial (universal) 16S rRNA gene via the Eub519/Univ907 primer set (Burgmann et al., 2011).

Bio-Rad iQ SYBR Green Supermix (Bio-Rad, Hercules, CA, USA) containing 50 U/ml iTaq DNA polymerase, 0.4 mM dNTPs, 100 mM KCl, 40 mM Tris-HCl, 6 mM MgCl2, 20 mM fluorescein, and stabilizers was used for all qPCR assays. The final volume of the reaction mix for each PCR and qPCR reaction was 20 µl, in which the DNA template was ~1 ng, and the primer concentrations were 0.2 µM. All assays were performed in triplicate. For each assay, triplicate standard series were generated by tenfold serial dilutions ($10^2$-$10^8$ gene copies/µl). All assays employed thermocycling conditions reported in the reference papers, and were performed on a Bio-Rad C1000 CFX96 Real-Time PCR system (Bio-Rad, Hercules, CA, USA).

#### S1.4 Comammox *amoA* cloning and sequencing

198 bp fragments of the comammox *amoA* gene were amplified, cloned, and sequenced from DNA extracts of the sludge sampled on day 407 in order to generate standards for qPCR and confirm specificity of the comammox primer set (Fowler et al., 2018). Triplicate PCR products were pooled and purified via gel electrophoresis using the Qiagen QiaQUICK Gel Extraction Kit

(Qiagen, Hilden, Germany). The purified PCR products were cloned using the TOPO® TA cloning® kits (Invitrogen/Thermo Fisher Scientific, Waltham, MA, USA) per the manufacturer's protocol. After blue/ white colony screening, three colonies were randomly selected and cultured overnight in LB medium containing 50 µg/ml ampicillin. The plasmids were isolated using the Invitrogen PureLink Quick Plasmid Miniprep Kit (Invitrogen/Thermo Fisher Scientific), and cloned inserts sequenced using the M13F-20 primer on the ABI 3730 automated Sanger sequencer at ACTG, Inc. (Wheeling, IL, USA). The acquired sequences were deposited in GenBank, and accession numbers can be found in SI Table S2.

**S1.5 16S rRNA, *amoA,* and *nxrB* gene PCR amplification, amplicon sequencing, analysis and phylogenetic inference**

16S rRNA and functional gene amplicon library preparations were performed using a two-step PCR protocol using the Fluidigm Biomark: Multiplex PCR Strategy. In the first round of PCR, each 20 uL reaction contained 10 µL of FailSafe PCR 2X PreMix F (Epicentre, Madison, WI), 0.63 units of Expand High Fidelity PCR Taq Enzyme (Sigma-Aldrich, St. Louis, MO), 0.4 µM of forward primer and reverse primer modified with Fluidigm common sequences at the 5' end of each primer, 1 µL of gDNA (approximately 100 ng) and the remaining volume molecular biology grade water. The V4-V5 region of 16S rDNA was amplified in duplicate from 27 samples collected over the course of reactor operation using the 515F-Y (5'-GTGYCAGCMGCCGCGGTAA-3') and 926R (5'-CCGYCAATTYMTTTRAGTTT-3') (Parada et al., 2016) primer set. Thermocycling conditions for the 515F-Y/926R primer set were 95°C for 5 minutes, then 28 cycles of 95°C for 30 seconds, 50°C for 45 seconds, and 68°C for 30 seconds, followed by a final extension of 68°C for 5 minutes. Amplification was checked for all samples (including those described below) via agarose gel electrophoresis.

To characterize overall *Nitrospira* community structure in this system, *Nitrospira nxrB* amplicon sequencing was performed on samples from day 407 of reactor operation using the 169f (5'-TACATGTGGTGGAACA) and 638R (5'-CGGTTCTGGTCRATCA) primer set (Pester et al., 2014). Thermocycling conditions were 95°C for 5 minutes, then 35 cycles of 95°C for 40 seconds, 56.2°C for 40 seconds, and 72°C for 90 seconds.

To assess community structure of comommox *Nitrospira*, comammox *amoA* amplicon sequencing was performed on samples from day 407 using the Ntsp-amoA 162F (GGATTTCTGGNTSGATTGGA) and Ntsp-amoA 359R (WAGTTNGACCACCASTACCA) primer set (Fowler et al., 2018). Thermocycling conditions were 94°C for 5 minutes, then 35 cycles of 94°C for 45 seconds, 56°C for 30 seconds, 72°C for 60 seconds, followed by a final extension of 72°C for 10 minutes. Amplification was checked for all samples on a 1% agarose gel.

Samples were then barcoded by sample via a second stage PCR amplification using Access Array Barcodes (Fluidigm, South San Francisco, CA)(Griffin and Wells, 2017). Each 20 uL PCR reaction consisted of 10 µL of FailSafe PCR 2X PreMix F, 0.63 units of Expand High Fidelity PCR Taq Enzyme, 2 µL of template from the first round of PCR, 4 µL of sample-specific barcode primers and the remaining volume molecular biology grade water. The conditions for the second round of PCR were 95°C for 5 minutes, then 8 cycles of 95°C for 30 seconds, 60°C for 30 seconds, and 68°C for 30 seconds. Agarose gel electrophoresis was run again after the second round of PCR to verify correct amplification. Amplicons were pooled and then purified using Ampure XP bead cleanup, checked for quantity and quality using Qubit fluorescence analysis and TapeStation2200, respectively, and then diluted to appropriate concentration for sequencing. Sequencing was performed on an Illumina Miseq sequencer (Illumina, San Diego, CA) using Illumina V2 (2x250 paired end) chemistry.

For all amplicon sequence analysis, the primers were trimmed and the pair-end reads were merged using Trimmomatic. The 16S rRNA gene amplicons were initially clustered into operational taxonomic units (OTUs) at 97% identity in VSEARCH (Rognes et al., 2016). After de novo chimera filtering in VSEARCH, OTU taxonomy was assigned using the Silva database (Release 128) using the Quantitative Insights Into Microbial Ecology (QIIME) (Quast et al., 2013) command align_seqs.py with the Muscle option. To identify biologically distinct amplicons at sub-OTU resolution, 16S rRNA gene amplicons that were taxonomically assigned to *Nitrospira* were reclassified into unique sequences (also called amplicon sequence variants or ASV) using DADA2 with default settings (Callahan et al., 2016). Functional gene amplicons (*nxrB* and *amoA*) were also classified with DADA2.

Phylogenetic analyses were conducted on *Nitrospira* 16S rRNA, *Nitrospira nxrB*, and comammox *amoA* gene amplicon sequences. The *Nitrospira* 16S rRNA genes (fragment length = 370 bp) were aligned in the ARB software package (Ludwig, 2004), and the maximum-likelihood phylogenetic tree was constructed with RAxML (Stamatakis, 2014). The *nxrB* (fragment length = 453 bp) and *amoA* (fragment length = 173 bp) gene sequences were aligned with MAFFT (Katoh and Standley, 2013), and the maximum-likelihood phylogenetic trees were constructed with FastTreeMP (Price et al., 2010). All the trees were visualized in iTOL (Letunic and Bork, 2016). The *nxrB* gene from *Nitrobacter hamburgensis* X14 and the *amoA* gene from *Nitrosococcus watsonii* C-113 were applied as outgroups in the *nxrB* and *amoA* trees, respectively. The 16S rRNA gene from *Nitrobacter winogradskyi* ATCC 14123 was applied as outgroup in the Nitrospira 16S rRNA gene tree in Figure S3; and the 16S rRNA gene from *Escherichia Coli* DE147 was applied as the outgroup in the phylogenetic tree for FISH probe evaluation in Figure S5. The bootstrap value indicating the reliability of each split in the tree were all represented as gray circles in the inner nodes of the tree. FastTreeMP applies the Shimodaira-Hasegawa test and uses 1,000 resamples to generate the bootstrap value of each split, and RAxML applies 1,000 bootstrap runs and the GTR substitution model to generate the bootstrap value of each split in the tree.

**S1.6 Fluorescent *in situ* Hybridization**

Fluorescent *in situ* hybridization (FISH) was performed to estimate the relative abundance of canonical *Nitrospira*, canonical ammonia oxidizing bacteria (AOB), a subset of canonical *Nitrospira* that includes comammox, and total Eubacteria (see Table S1 for probe sequences and corresponding references). Reactor biomass samples were chemically fixed in a 4% formaldehyde solution for two hours at 4°C followed by storage at -20°C in a 1:1 solution of phosphate buffered saline (PBS) and ethanol. Fixed biomass samples were diluted 6-fold in PBS and homogenized on ice in a Potter-Elvehjem tissue grinder (Kontes, Model# 886000-0020). 10 µl of the diluted homogenized samples were transferred to polytetrafluoroethylene (PTFE) printed 8-well slides and incubated at 60 °C for 1 hour to improve adhesion of the dehydrated biomass to the slide.

Samples underwent a dual-staining hybridization following protocols adapted from Nielsen et al. (2009). For the "*Nitrospira*-AOB-Ntspa476" (see section S2. for discussion on probe Ntspa476 specificity) treatment, samples were first hybridized for four hours in a 35% formamide v/v hybridization buffer containing canonical *Nitrospira* probes (Ntspa662, Ntspa712; 6-FAM fluorophore) at concentrations of 0.83 µM and canonical Betaproteobacterial ammonia oxidizing bacteria probes (NEU, Nso1225, Cluster6a192; Cy3 fluorophore) at concentrations of 0.5 µM. Samples were then hybridized for four hours in a 20% formamide v/v hybridization buffer containing 0.5 µM Ntspa476 (Cy5 fluorophore), which was originally developed for the detection of the putative comammox organisms *Ca.* N. nitrosa and *Ca.* N. Nitrificans (van Kessel et al.,

2015). For the "Eub-AOB-Ntspa476" treatment, the canonical *Nitrospira* probes were replaced with Eub338 I, Eub338 II, and Eub338 III ("Eub Mix"; 6-FAM fluorophore) at concentrations of 0.83 μM. The Eub mix was also added to the 20% formamide hybridization at concentrations of 0.83 μM. All probe mixes included equimolar concentrations of the recommended competitor oligonucleotides listed in Table S1. Each sample was separately stained with the nonsense probe (Non-EUB338) tagged with 6-FAM, Cy3, and Cy5 fluorophores as a negative control for non-specific binding.

Hybridized cryosections were counterstained with DAPI at a concentration of 1 μg/mL in a glycerol based antifadent mounting media (Citifluor AF1). Image stacks were acquired on an inverted confocal laser scanning microscope (Model TCS SP5, Leica Microsystems) equipped with an oil immersion 63× (1.44 NA) objective at a lateral resolution of 0.48 μm and an axial step size of 1 μm. DAPI, 6-FAM, Cy3, and Cy5 fluorophores were excited sequentially with 405nm, 488nm, 561nm, and 633nm laser lines, respectively. Ten fields of view were collected for each sample treatment.

Ratios of Ntspa476/AOB/*Nitrospira* and Ntspa476/AOB/total Eubacteria were calculated in the Fiji software package (Schindelin et al., 2012). Binary thresholds for each image channel were independently selected by a panel of three image analysts to distinguish hybridized biomass from background fluorescence. A given microbial quantity (e.g. AOB, *Nitrospira*) was calculated as the average thresholded pixel count in a given channel across ten fields of view averaged across the corresponding values reported by the three image analysts.

**S2. Analysis of FISH probe Ntspa476 specificity**

The specificity and coverage of the putative comammox FISH probe Ntspa476 (van Kessel et al., 2015) and the *Nitrospira* FISH probe mix Ntspa662 and Ntspa712 (Daims et al., 2001) were evaluated by phylogenetic analysis on database-derived 16S rRNA gene sequences that could predictably hybridize with these probes. For this purpose, the Silva 132 SSURef NR99 database (695,171 sequences in total) was downloaded and imported into the ARB software (Ludwig, 2004). Probe_server.arb was built on the whole SSU dataset. In this *in silico* analysis, we assumed database sequences with 0 mismatch to the FISH probe mix would hybridize with the probe. 726 sequences that had no nucleotide mismatches to either the Ntspa662 or the Ntspa712 probes were exported as sequences predicted to hybridize with the *Nitrospira*-specific FISH probe mix (Ntspa662 and Ntspa712). We also exported the 44 sequences that had no mismatches to the Ntspa476 FISH probe; 43 of these were a subset of the previous 726 sequences predicted to also hybridize to the *Nitrospira* FISH probe mix (Ntspa662 and Ntspa712). All exported 16S rRNA gene sequences, together with the 5 dominant *Nitrospira* 16S rRNA gene ASV recovered via amplicon sequencing in this study, were aligned in ARB and the phylogenetic tree shown in Figure S5 was generated with the RAxML software applying the GTR substitution model (Stamatakis, 2014).

43 out of the 44 sequences that we predicted would hybridize with Ntspa476 were also predicted to hybridize with the *Nitrospira* FISH probe mix (Ntspa662 and Ntspa712), and these sequences are shown in light magenta in Figure S5. The 1 sequence predicted to hybridize to Ntspa476 but not to Ntspa662 or Ntsap712 is shown in dark magenta. The 673 (i.e. 726 – 43 = 673) 16S rRNA genes that hybridize only with the *Nitrospira* FISH probe mix (Ntspa662 and Ntspa712) but not with Ntspa476 are shown in blue in Figure S5. As is indicated in Figure S5, the specificity of the *Nitrospira* FISH probe mix (Ntspa662 and Ntspa712) is reasonably good, with minimal hybridizing sequences lying beyond the phylum of Nitrospirae. Of 726 total 16S rRNA sequences predicted to hybridize to either Ntspa662 or Ntspa712, only 19 (2.6%) did not affiliate

with the Nitrospirae phylum (3 sequences from the phylum of Planctomycetes, 11 sequences from the phylum of Chloroflexi, 2 sequences from the phylum of Proteobacteria, and 3 sequences from the phylum of BRC1). All 44 sequences predicted to hybridize with the FISH probe Ntspa476 are in the genus *Nitrospira*, and 98% (43/44) of them are in *Nitrospira* lineage II. However, the resolution of FISH probe Ntspa476 in distinguishing the 16S rRNA genes between comammox and canonical NOBs is uncertain, based on the fact that gene sequences from known lineage II canonical *Nitrospira* (e.g. *Nitrospira lenta* (NR_148573.1)) specific to the *Nitrospira* probe mix but not to Ntspa476 were interspersed with 16S rRNA gene sequences predicted to hybridize to Ntspa476. Taken together, this analysis suggests that probe Ntspa476 is likely not capable of reliably resolving comammox from conventional nitrite-oxidizing *Nitrospira*, but is capable of specifically targeting a subset of lineage II *Nitrospira* that includes known comammox *Nitrospira*.

Regarding the dominant *Nitrospira* 16S rRNA gene ASV (black text, Figure S5) identified in the present study, the three sequences in OTU4 that declined over time during reactor operation (see Figure 3) affiliated to *Nitrospira* lineage I. No *Nitrospira* 16S rRNA genes in lineage I are predicted to hybridize with probe Ntspa476. The two ASV in OTU2, i.e., 'NU_Nitrospira_16S_Rep_Seq1' and 'NU_Nitrospira_16S_Rep_Seq2', affiliated to *Nitrospira* lineage II, with the former phylogenetically closer to *Ca.* Nitrospira nitrosa and the latter much more abundant (as in shown in the heatmap of Figure S5). Given the high abundance of taxa hybridized to probe Ntspa476 in qFISH quantification data (comprising 75% of all *Nitrospira* by day 398, see Figure 6), it is reasonable to suggest that both of the 16S rRNA genes in OTU2 were hybridized by the comammox FISH probe, but it is uncertain whether the more abundant one – 'NU_Nitrospira_16S_Rep_Seq2' – is, in fact, comammox *Nitrospira*.

**Table S1.** 16S rRNA-targeted oligonucleotide FISH probes used in this study.

| Probe | Sequence (5' to 3') | FA (%) | Specificity | Reference |
|---|---|---|---|---|
| NEU | CCC CTC TGC TGC ACT CTA | 35 | Most halophilic and halotolerant *Nitrosomonas* spp. | (Wagner et al., 1995) |
| CTE (NEU Competitor) | TTC CAT CCC CCT CTG CCG | 35 | Unlabeled together with NEU *Comamonas* spp., *Acidovorax* spp., *Hydrogenophaga* spp., *Aquaspirillum* spp. | (Wagner et al., 1995) |
| Nso1225 | CGC CAT TGT ATT ACG TGT GA | 35 | Ammonia-oxidizing β-proteobacteria | (Mobarry et al., 1996) |
| Cluster6a192 | CTT TCG ATC CCC TAC TTT CC | 35 | *Nitrosomonas oligotropha* lineage (Cluster 6a) | (Adamczyk et al., 2003) |
| Cluster6a 192 Competitor | CTT TCG ATC CCC TGC TTT CC | 35 | Competitor for Cluster6a192 | (Adamczyk et al., 2003) |
| Ntspa662 | GGA ATT CCG CGC TCC TCT | 35 | Genus *Nitrospira* | (Daims et al., 2001) |
| Ntspa662 Competitor | GGA ATT CCG CTC TCC TCT | 35 | Competitor for Ntspa662 | (Daims et al., 2001) |
| Ntspa712 | CGC CTT CGC CAC CGG CCT TCC | 35 | Most members of the phylum Nitrospirae | (Daims et al., 2001) |
| Ntspa712 Competitor | CGC CTT CGC CAC CGG TGT TCC | 35 | Competitor for Ntspa712 | (Daims et al., 2001) |
| Ntspa476 | CTG CAG GTA CCG TCC GAA | 20 | *Ca.* N. nitrosa, *Ca* N. nitrificans | (van Kessel et al., 2015) |
| Ntspa476 Competitor | CTG GAG GTA CCG TCC GAA | 20 | Competitor to Ntspa476 | (van Kessel et al., 2015) |
| Eub338 | GCT GCC TCC CGT AGG AGT | 20, 35 | Most Bacteria | (Amann et al., 1990) |
| Eub338 II | GCA GCC ACC CGT AGG TGT | 20, 35 | Planctomycetales | (Daims et al., 1999) |
| Eub338 III | GCT GCC ACC CGT AGG TGT | 20, 35 | Verrucomicrobiales | (Daims et al., 1999) |
| NON Eub | ACT CCT ACG GGA GGC AGC | 20, 35 | Negative control for nonspecific binding | (Wallner et al., 1993) |

**Table S2.** GenBank accession numbers for *Nitrospira nxrB* and comammox *amoA* amplicon sequence variants, comammox *amoA* clone sequences, and raw sequencing data.

| Amplicon sequence variant ID | GenBank accession number | Amplicon sequence variant ID | GenBank accession number |
|---|---|---|---|
| NU_nxrB_0 | MH587129 | NU_nxrB_40 | MH587152 |
| NU_nxrB_1 | MH587130 | NU_nxrB_42 | MH587153 |
| NU_nxrB_2 | MH587131 | NU_nxrB_47 | MH587154 |
| NU_nxrB_3 | MH587132 | NU_nxrB_54 | MH587155 |
| NU_nxrB_4 | MH587133 | NU_nxrB_59 | MH587156 |
| NU_nxrB_6 | MH587134 | NU_nxrB_82 | MH587157 |
| NU_nxrB_7 | MH587135 | NU_nxrB_86 | MH587158 |
| NU_nxrB_8 | MH587136 | NU_Commmox_amoA4 | MH587119 |
| NU_nxrB_9 | MH587137 | NU_Commmox_amoA5 | MH587120 |
| NU_nxrB_12 | MH587138 | NU_Commmox_amoA3 | MH587121 |
| NU_nxrB_13 | MH587139 | NU_Commmox_amoA7 | MH587122 |
| NU_nxrB_17 | MH587140 | NU_Commmox_amoA2 | MH587123 |
| NU_nxrB_19 | MH587141 | NU_Commmox_amoA8 | MH587124 |
| NU_nxrB_20 | MH587142 | NU_Commmox_amoA1 | MH587125 |
| NU_nxrB_22 | MH587143 | NU_Commmox_amoA_clone1 | MH587126 |
| NU_nxrB_23 | MH587144 | NU_Commmox_amoA_clone3 | MH587127 |
| NU_nxrB_24 | MH587145 | NU_Commmox_amoA_clone2 | MH587128 |
| NU_nxrB_25 | MH587146 | NU_Nitrospira_16S_rep_seq2 | MH587159 |
| NU_nxrB_26 | MH587147 | NU_Nitrospira_16S_rep_seq5 | MH587160 |
| NU_nxrB_27 | MH587148 | NU_Nitrospira_16S_rep_seq13 | MH587161 |
| NU_nxrB_28 | MH587149 | NU_Nitrospira_16S_rep_seq14 | MH587162 |
| NU_nxrB_29 | MH587150 | NU_Nitrospira_16S_rep_seq117 | MH587163 |
| NU_nxrB_30 | MH587151 | 16S rRNA/*amoA*/*nxrB* raw data | PRJNA480047 |

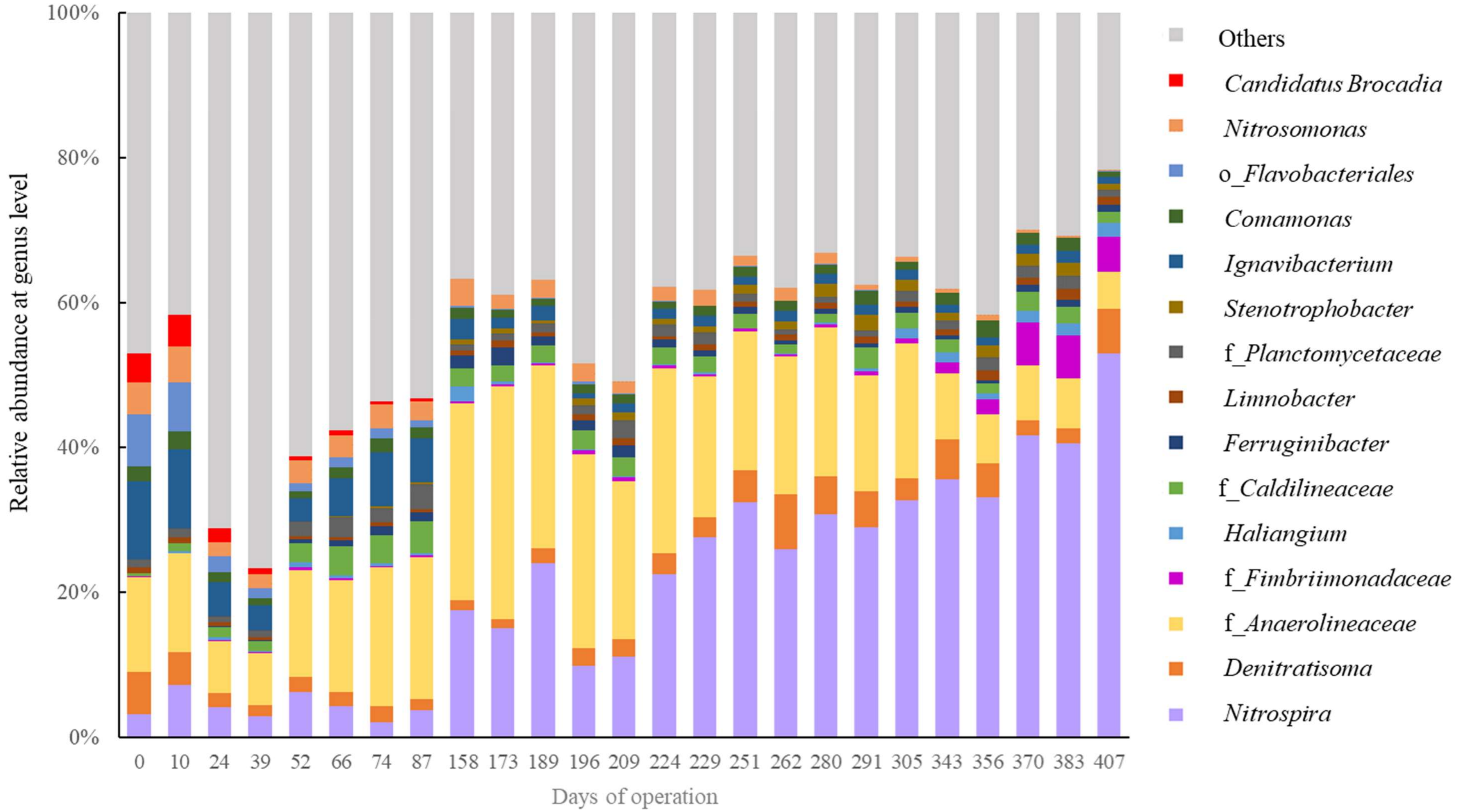


**Figure S1.** Relative abundance of the 15 most abundant bacterial genera at each time point according to 16S rRNA gene sequencing data. Sampling time point is shown on the x axis. The OTUs that were unclassified at the genus level are presented with the corresponding lowest annotable taxonomy names; ‘f_’ indicate ‘Family’, and ‘o_’ indicates ‘Order.’

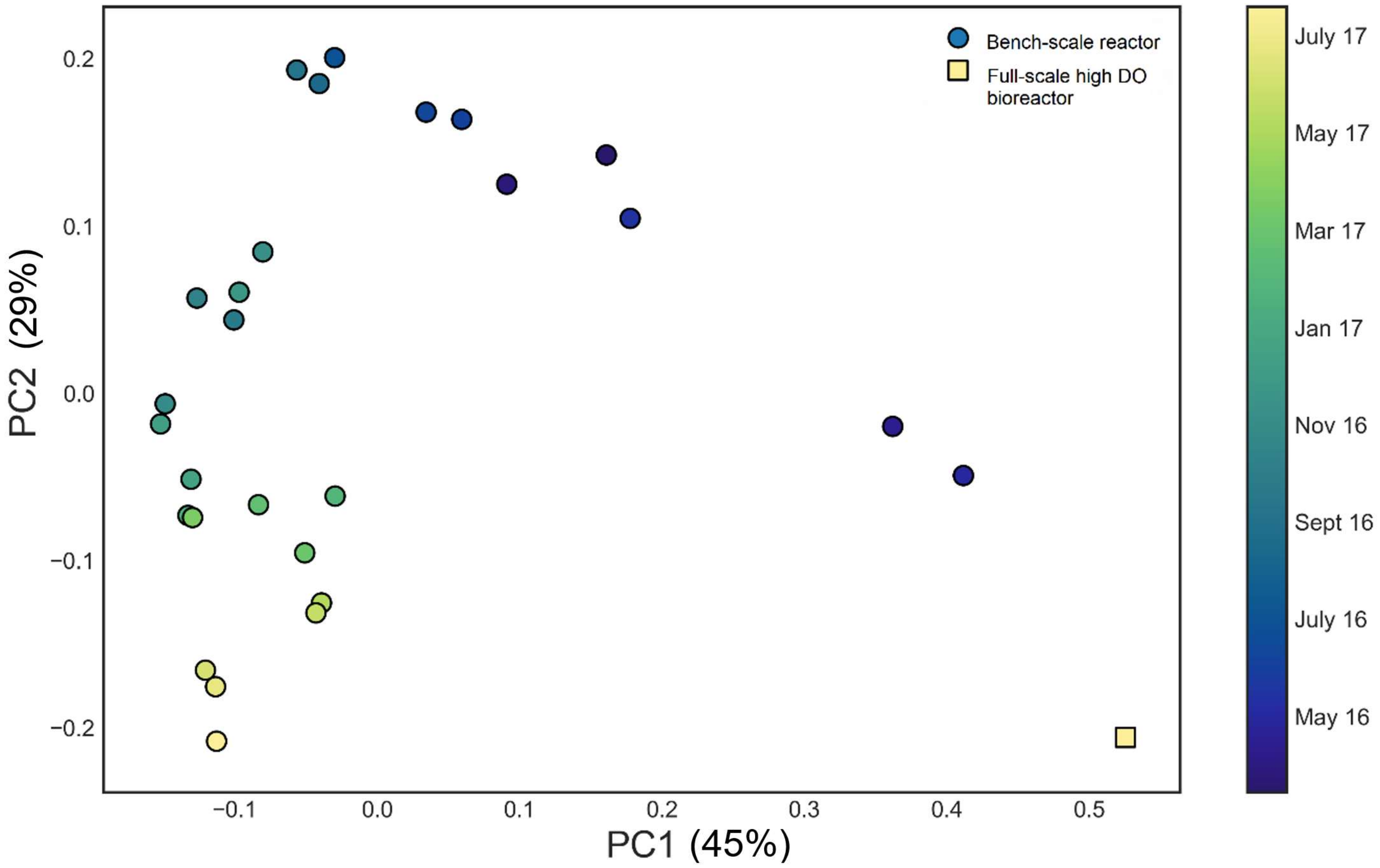


**Figure S2**. Principle Coordinate Analysis of weighted unifrac distances between samples, based on 16S rRNA gene amplicon sequencing analyses. Samples from the low DO bench-scale reactor are indicated by circles, and a single sample for comparison in the parallel full-scale high DO nitrifying activated sludge bioreactor is indicated by a square. Samples are color coded by time of sampling.

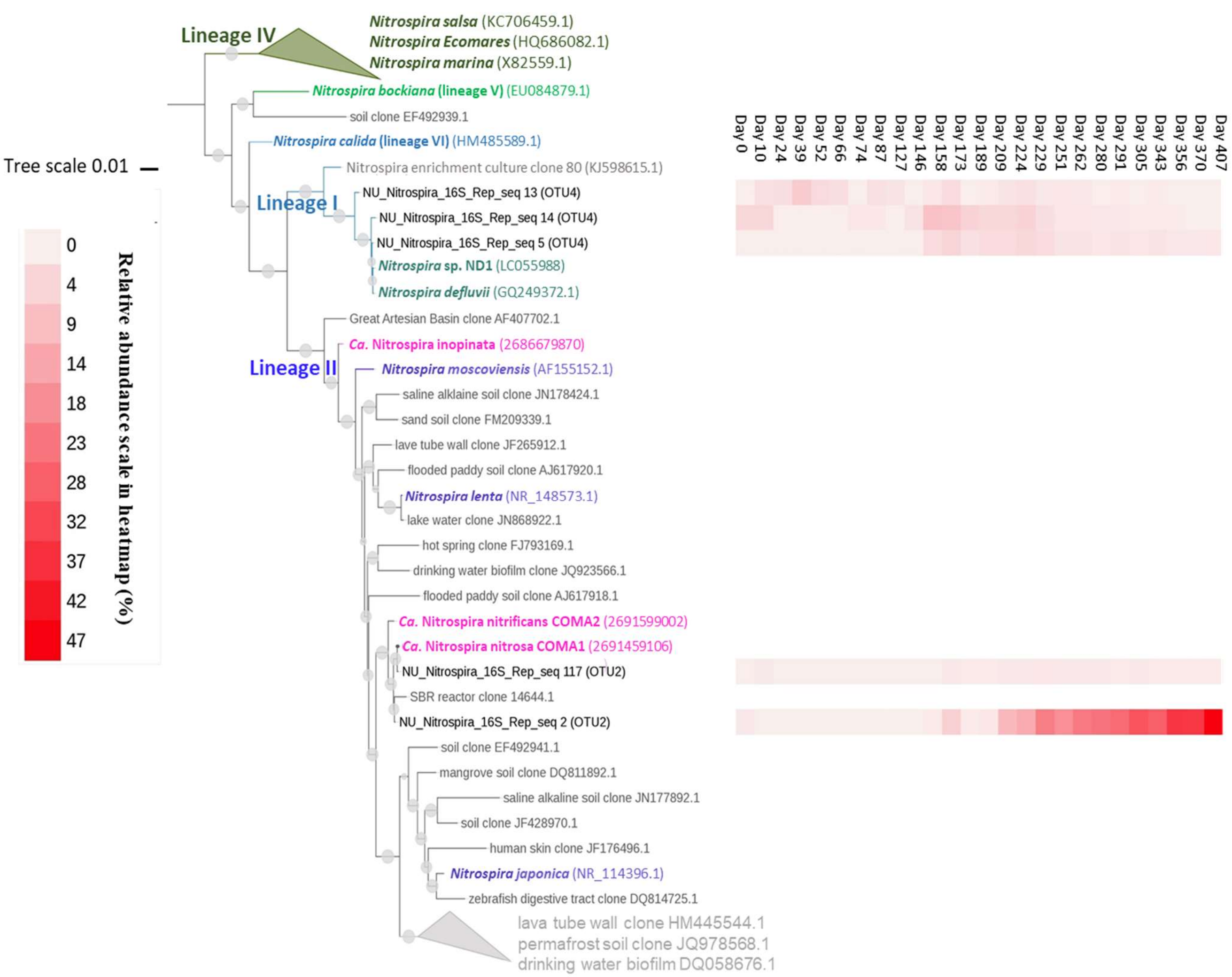


**Figure S3.** Maximum-likelihood phylogenetic tree of the 5 *Nitrospira* 16S rRNA gene amplicon sequence variants (ASV) (370 bp fragment) in the two dominant OTUs (OTU2 and OTU4, clustered to 97% identity). The heatmap shows the abundance of each of the 5 ASV against the total number of the 16S rRNA gene reads.

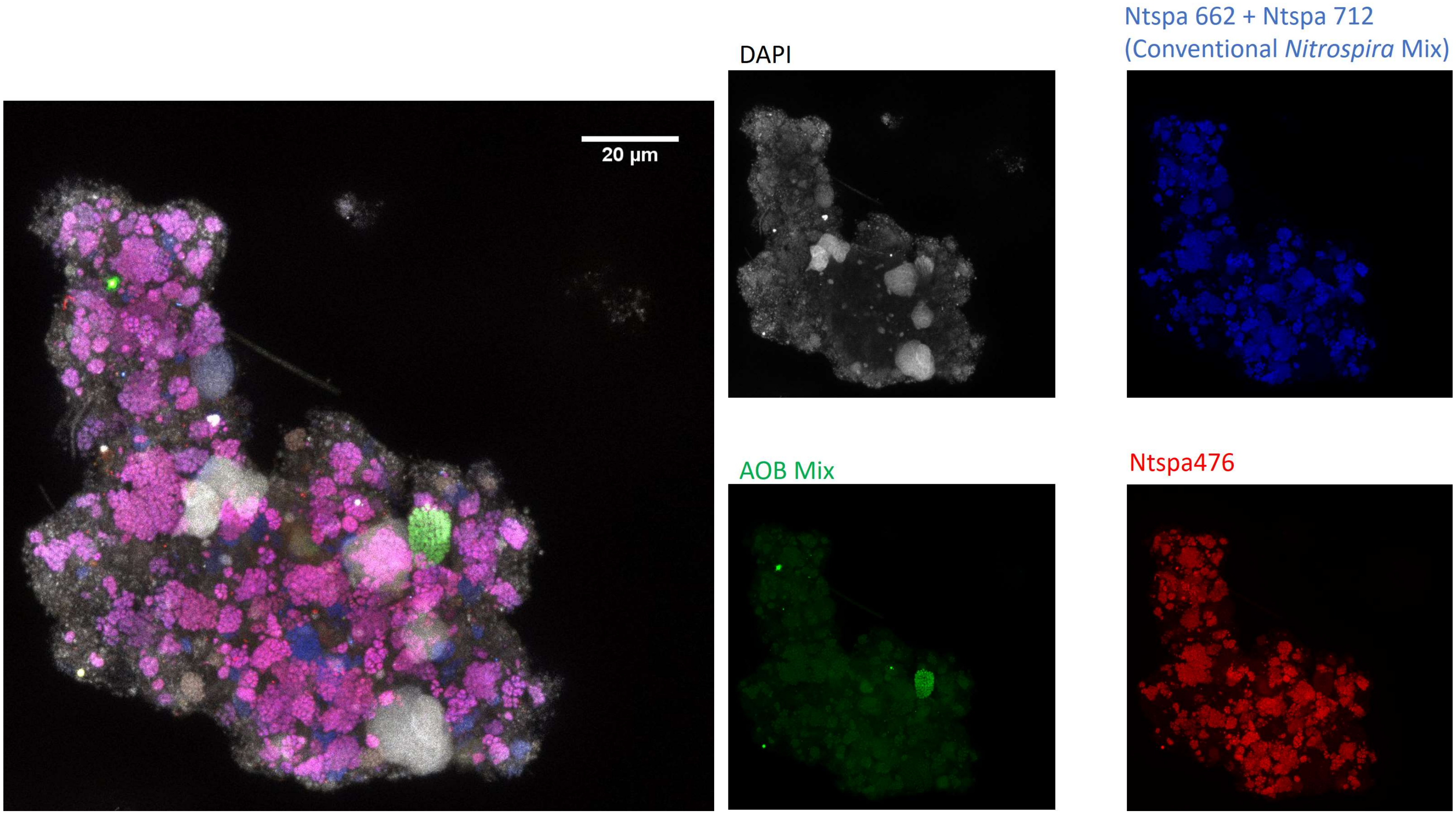


**Figure S4**. Representative FISH micrograph of showing co-aggregation of *Nitrospira*, *Nitrosomonas* and Ntspa476-hybridized *Nitrospira* in the low DO bench scale nitrification reactor consortium on day 407. DAPI counterstaining is in grey, and probes specific for *Nitrospira* (Ntspa662 and Ntspa712, blue), a subset of *Nitrospira* that includes known comammox (Ntspa476, red; red + blue = magenta), and *Nitrosomonas* (AOB mix: NEU, Nso1225, Cluster6a192; green)

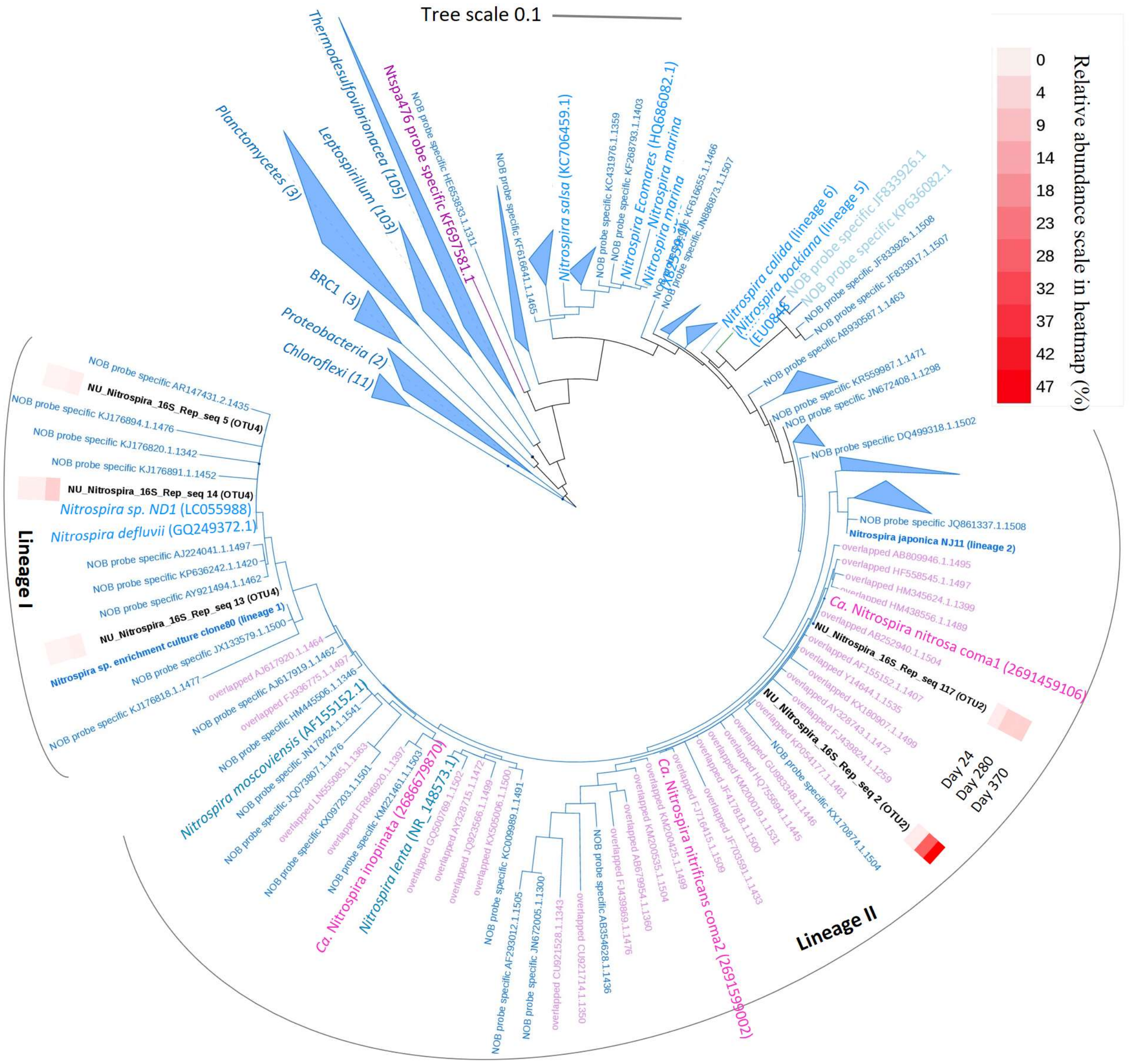


**Figure S5.** Maximum-likelihood phylogenetic tree of the 16S rRNA genes that could be targeted by FISH probe Ntspa476 (originally designed to target specific comammox taxa) and *Nitrospira* FISH probe mix (Ntspa662 and Ntspa712). Magenta indicates sequences predicted to hybridize with both Ntspa476 and the *Nitrospira* probe mix; blue indicates sequences predicted to hybridize with only the *Nitrospira* probe mix; pink indicates the 16S rRNA genes from three comammox genomes; and black indicates the five *Nitrospira* 16S rRNA gene ASV that were dominant in the reactor consortium of this study, with the relative abundance of each indicated in the heatmap. Numbers in parentheses after taxonomy names indicate the number of 16S rRNA genes that are predicted to hybridize with the *Nitrospira* FISH probes (Ntspa662 or Ntspa712) in the corresponding taxa. The heatmap indicates the abundance of each the *Nitrospira* 16S rRNA gene ASV against the total number of the 16S rRNA gene reads.
